# geneBasis: an iterative approach for unsupervised selection of targeted gene panels from scRNA-seq

**DOI:** 10.1101/2021.08.10.455720

**Authors:** Alsu Missarova, Jaison Jain, Andrew Butler, Shila Ghazanfar, Tim Stuart, Maigan Brusko, Clive Wasserfall, Harry Nick, Todd Brusko, Mark Atkinson, Rahul Satija, John Marioni

## Abstract

The problem of selecting targeted gene panels that capture maximum variability encoded in scRNA-sequencing data has become of great practical importance. scRNA-seq datasets are increasingly being used to identify gene panels that can be probed using alternative molecular technologies, such as spatial transcriptomics. In this context, the number of genes that can be probed is an important limiting factor, so choosing the best subset of genes is vital. Existing methods for this task are limited by either a reliance on pre-existing cell type labels or by difficulties in identifying markers of rare cell types. We resolve this by introducing an iterative approach, geneBasis, for selecting an optimal gene panel, where each newly added gene captures the maximum distance between the true manifold and the manifold constructed using the currently selected gene panel. We demonstrate, using a variety of metrics and diverse datasets, that our approach outperforms existing strategies, and can not only resolve cell types but also more subtle cell state differences. Our approach is available as an open source, easy-to-use, documented R package (https://github.com/MarioniLab/geneBasisR).

## Introduction

Single-cell RNA sequencing (scRNA-seq) is a fundamental approach for studying transcriptional heterogeneity within individual tissues, organs and organisms (reviewed in ^1^). A key step in the analysis of scRNA-seq data is the selection of a set of representative features, typically a subset of genes, that capture variability in the data and that can be used in downstream analysis. Established approaches for feature selection leverage quantitative per gene metrics that aim to identify genes that display more variability than expected by chance across the population of cells under study. Commonly used methods for detecting highly variable genes (HVG) utilise the relationship between mean and standard deviation of expression levels (reviewed in ^2^), GiniClust leverages Gini indices ^3^, and M3Drop performs dropout-based feature selection ^4^. The number of selected genes is typically dependent on a user-defined threshold, but ordinarily is on the order of one to a few thousand genes ^2,5^. A recently developed approach, scPNMF, further addresses the gene complexity problem by leveraging a Non-Negative Matrix Factorisation (NMF)-representation of scRNA-seq, with selected features being suggested to represent interesting biological variability in the data ^6^. scPNMF relies on the chosen dimension for the NMF-representation, and also does not directly compare informativeness between different factors, thus impeding the ability to compare the importance (i.e. gene weights) between different factors.

Framing the problem as searching for a *fixed* number of informative genes has recently gained practical relevance. Specifically, recent technological advances have given rise to multiple approaches where gene expression can be probed at single-cell resolution within a cell’s spatial context, thus enabling the relationship between individual cells and cell types to be studied (reviewed in ^7–9^). One branch of spatial transcriptomics approaches builds on the concept of RNA Fluorescence In Situ Hybridisation (FISH), where probes are used to assay the number of molecules of a given gene present in a single cell ^10^. Newly developed approaches, such as seqFISH ^11–14^, MERFISH ^15–17^ and DARTFISH ^18^ scale up classical RNA FISH strategies, allowing the expression of dozens to hundreds of genes to be assayed in parallel. The single-molecule resolution of these methods enables quantification of the location of individual RNA molecules inside a single cell, which has important implications for studying developmental, neuronal and immune biology, where cell-cell interactions are highly informative. However, an important limitation of such methods is that the number of genes that can be probed (gene panel) is limited (typically in the low hundreds) and this gene panel needs to be selected prior to the experiment taking place. Additionally, carefully selected targeted gene panels are relevant for CRISPR-seq screens ^19–21^ and, more recently, targeted single-cell gene expression assays ^22^ have been developed, which improves capture efficiency and reduces experimental cost.

Current strategies that address the problem leverage prior knowledge regarding the relevance of genes for the tissue of interest as well as using unsupervised marker selection, where matched scRNA-seq data is used to find genes of interest. If the goal is to identify genes that characterise clusters present in scRNA-seq data, existing marker selection methods can be applied, such as one-vs-all methods testing for genes that exhibit differential expression between clusters ^23–26^, ranking genes by correlation with the normalized boolean vectors corresponding to the clusters ^27^ or random forest classification algorithms ^28,29^. In addition to one-vs-all methods, the scGeneFit approach selects marker genes that jointly optimize cell type recovery using a label-aware compressive classification method, and returns the optimal set of markers given a user-defined panel size ^30^.

The main drawback of these approaches is that they fundamentally rely on pre-determined cell type annotation, thus relying not only on the clustering algorithm but also that the correct granularity of clustering has been employed. By design, genes associated with intra-celltype variation or genes that vary in expression over a subset of clustered cell types will not be captured by such approaches. While the latter type of signal is generally deemed uninteresting, in practice this will depend on the system and question in mind. For example, tracing stress response and DNA damage will be relevant when analyzing cancerous samples, and understanding proliferation rates and cell cycle state is important in numerous biological contexts.

These fundamental limitations of most existing marker selection methods can be resolved by deploying unsupervised selection methods that aim to capture features (genes) that describe all sources of variability present in scRNA-seq data, both within and across cell types. One recently developed method, SCMER, addresses this problem by aiming to find a set of genes that preserve the overall manifold structure of scRNA-seq data ^31^. While novel and algorithmically impressive, SCMER has limitations that hinder its utility in practice. First, SCMER strives to find a selection that preserves the whole manifold and returns a scalar estimate for how well the manifold is preserved in the form of a KL-divergence between the ‘true’ and the ‘selected’ manifolds. However, this global score reflects the overall preservation of manifold structure and may not adequately weight or represent the local preservation of rare cell types. Additionally, to deal with batch effects, SCMER finds common sources of variation across most of the batches, which will be suboptimal for cell types that are present in only a few batches. This is not an uncommon case for datasets collected from different donors and different tissue sites.

To overcome these limitations, we have developed a novel cluster-free, batch-aware and flexible approach, **geneBasis**, which takes scRNA-seq data and the number of genes to be selected and returns a ranked gene panel of the designated size. Importantly, we provide a comprehensive set of metrics – at multiple levels of granularity - that can be used to assess the completeness of the suggested panel. To incorporate expert knowledge, we allow users to pre-select genes of interest. Additionally, by avoiding the use of predefined clusters, geneBasis allows discovery of genes that underpin transcriptional programs within any selected groups of cells (e.g. cell types). We demonstrate the power of our approach by applying it in a variety of biological contexts and compare its performance to existing state-of-the-art strategies.

## Results

### An algorithm for gene selection

For a given scRNA-seq dataset, we represent transcriptional similarities between cells using k-nearest neighbour (k-NN) graphs (Methods). A k-NN graph that is constructed using the entire transcriptome represents the ‘true’ relationships between cells as manifested by the similarities in expression levels of individual genes between cells and their ‘true’ neighbours. Subsequently we aim to find a selection of genes that yields a k-NN graph similar to the ‘true’ graph and is capable of predicting the true expression of any gene by using each cell’s neighbours.

Specifically, at each iteration (i.e. given the currently selected gene panel) we compare the graph constructed using the entire transcriptome (‘true’ k-NN graph) with a graph constructed using the current ‘selection’ (‘selection’ graph). At a single gene level, we assess how well we can predict a gene’s expression levels across cells, using the cells neighbours in the ‘selection’ graph compared to the ‘true’ graph, and select the gene that shows the biggest discrepancy between the two graphs (**Fig. 1**). More precisely, for each gene, in the ‘true’ graph we compute, across cells, the Minkowski distance between a gene’s expression in a given cell and its average expression in that cell’s k nearest neighbours. Intuitively, this provides a baseline measure for how well a gene’s expression can be predicted by its nearest neighbours in a best-case scenario. Similarly, we compute the Minkowski distance for each gene using the ‘selection’ graph. We then compute, for each gene, the difference in the distances between the ‘true’ and ‘selection’ graph and add the gene with the largest difference in distance to the selected set. This process is then repeated until the desired number of genes has been chosen.

**Figure 1.**
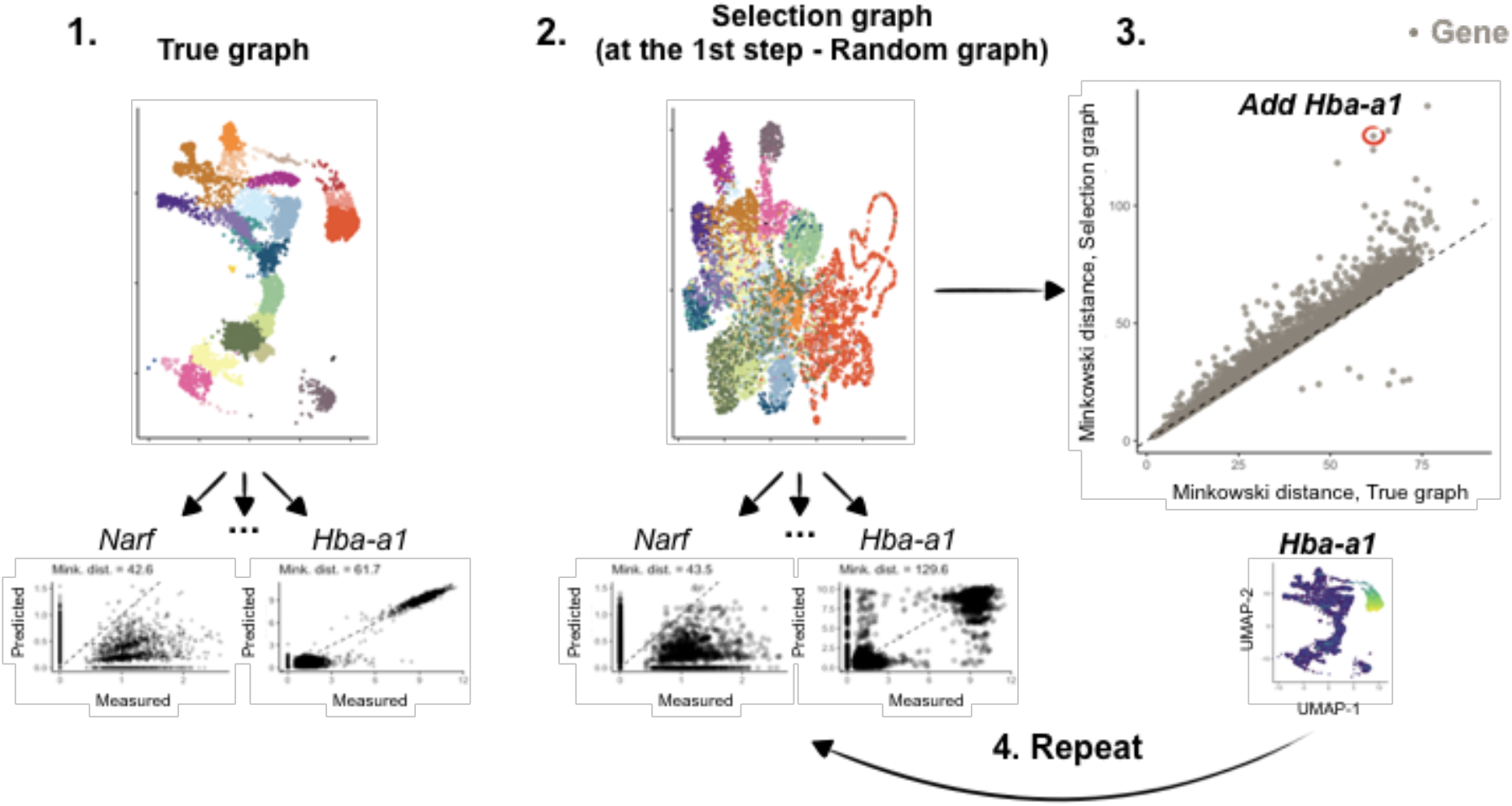
Schematic overview of geneBasis. Below we describe the steps of the algorithm. 1. ‘True’ k-NN graph is constructed, which represents the ‘ground truth’ relationship between cells. Colours correspond to celltypes for visualisation but are not used by the algorithm. For each gene (*Narf* and *Hba-a1* are used as examples) we calculate the Minkowski distance between two vectors: the first vector corresponds to log-normalised expression for each cell (referred to as measured); the second vector corresponds to the average log-normalised expressions across the first K neighbours for each cell (referred to as predicted). 2. For the current selection the ‘Selection’ graph is constructed to represent the relationship between cells using currently selected genes. As in 1., for each gene we calculate the Minkowski distance between measured and predicted counts. 3. The Minkowski distances are compared for each gene, and the gene with the biggest difference between Selected and True graphs (in the schematic - *Hba-a1*) is added to the current selection. 4. Steps 2-3 are repeated until the number of selected genes reaches the desired value.

An important limitation of existing methods is the inability to properly assess how complete and informative the designed gene panel is. To address this, we derived three metrics that evaluate gene panels at different levels of granularity: cell type, cell and gene level. The first metric explores whether cell types can be assigned in a specific and sensitive manner. Specifically, while geneBasis does not rely on clustering, for scRNA-seq datasets where annotations already exist, they can be exploited. To do so, we construct a k-NN graph for all cells using the selected gene panel, and compare for each cell, its (pre-determined) cell type label with the (pre-determined) labels assigned to its nearest neighbours (Methods). This allows construction of a confusion matrix that provides insight into whether cells from the same type are grouped together in the graph constructed using the selected panel.

As a second metric, which does not rely on cell type annotation, we focused on examining how well an individual cell’s neighbourhood is preserved. For each cell, we compute its nearest neighbours in the ‘true’ and ‘selected’ graph. Subsequently, in the ‘true’ graph space, we compare the distance between each cell and these two sets of nearest neighbours (**Supp. Fig. 1**). Intuitively, when the set of neighbours is identical in both the ‘true’ and ‘selection’ graph, this metric will output a score of 1, whilst a score close to 0 indicates that a cell’s neighbours in the ‘selection’ graph are randomly distributed across the ‘true’ manifold.

Finally, a third metric for assessing the quality of the selected gene panels focuses on individual genes (Methods). For each gene, we compute the correlation (across cells) between its log-normalised expression values and its average log-normalised expression values across K nearest neighbours in a graph. We compute correlations for the ‘true graph’ and for the ‘selected graph’, and use the ratio (‘selected’/’true’) as the final gene prediction score.

### geneBasis allows recovery of local and global variability

To evaluate whether geneBasis efficiently recovers a selection of genes that preserves both local and global structure, we applied geneBasis, as well as SCMER and scGeneFit (Methods), to four diverse systems representing challenges that frequently arise in scRNA-seq datasets:

1. A study of mouse embryogenesis at embryonic day (E)8.5, coinciding with early organogenesis ^32^. This dataset contains both abundant and rare cell types as well as multiple differential trajectories.
2. A newly-generated dataset profiling cells from the adult human spleen. As part of the lymphatic system, the spleen contains transcriptionally similar immune cell types that are not straightforward to resolve with a limited number of genes.
3. Multiple independent studies of the adult human pancreas dataset, consisting of multiple batches from various donors and experiments ^33–38^.
4. Transcriptional profiling of melanoma samples from 19 donors collected at different stages of disease progression ^39^.

Overall, using the metrics defined above, the gene panels chosen by geneBasis (**Supp. Table 1**) yield better performance than those chosen by SCMER (**Fig. 2A,B**). This is true both in terms of preserving cell neighbourhoods and in terms of the fraction of cells for which the neighbourhood was poorly predicted (**Fig. 2A, Supp. Fig. 3A**). The effect was most noticeable when a small number of genes were selected, with differences in performance decreasing as the number of genes selected increased (with the exception of the spleen, where geneBasis consistently outperforms SCMER). This is of practical relevance for designing gene panels for FISH-based experiments, where the number of selected genes can be an important limiting factor.

**Figure 2.**
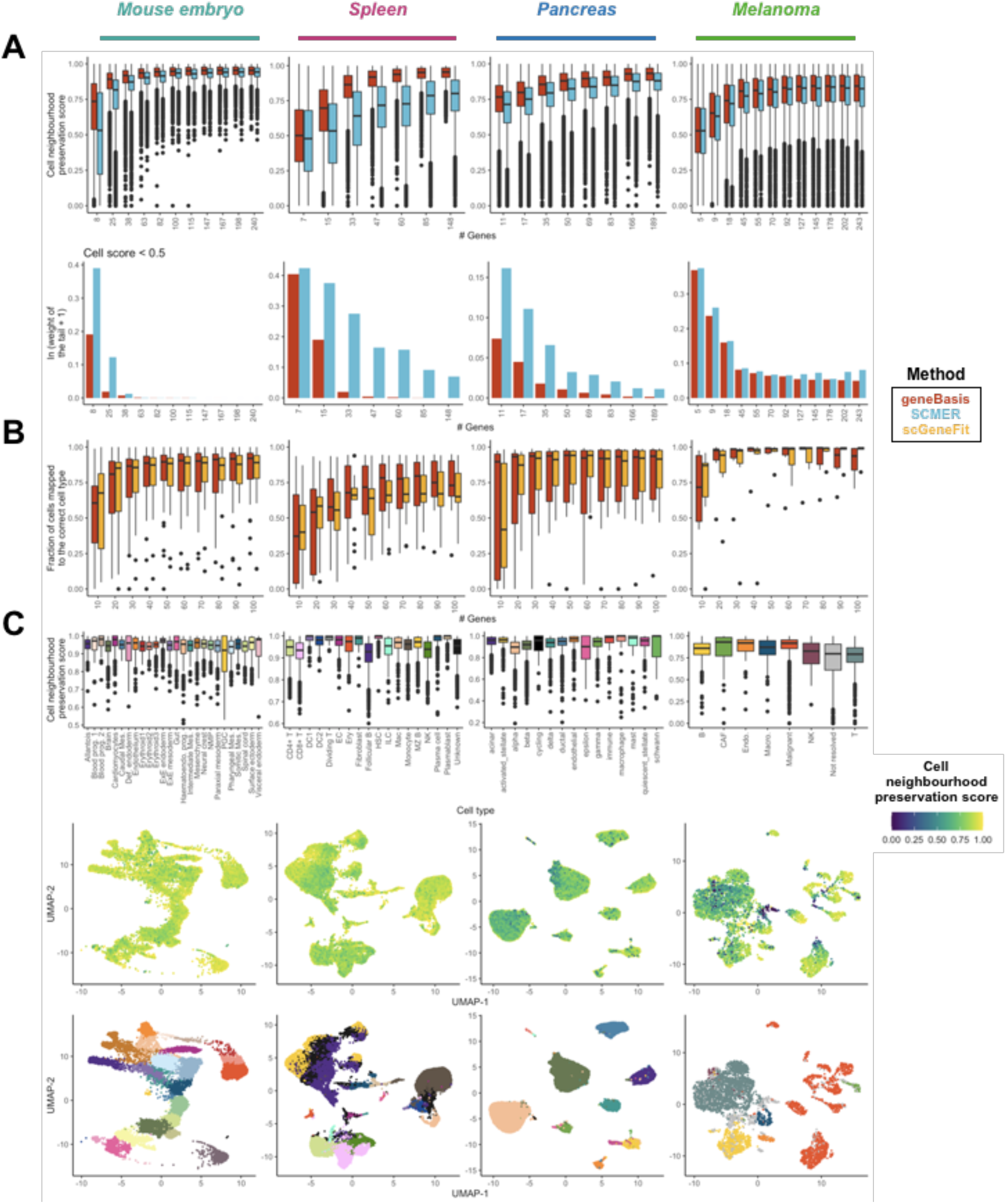
geneBasis preserves local and global structure. Each column represents the results for a different biological system. (A) Upper panel: The overall convergence of cell neighbourhood preservation distributions for geneBasis (in red) and SCMER (in blue) as a function of the number of selected genes. Lower panel: the weight of tails of the distributions (i.e. fraction of cells with neighborhood preservation score < 0.5). Values are rescaled as ln(x + 1). (B) The fraction of cells mapped to the correct cell type for geneBasis (in red) and scGeneFit (in orange) as a function of number of genes in the selection. (C) Upper panel: The distribution of neighbourhood preservation scores (calculated for the first 150 genes) per celltype. Middle panel: UMAP representation coloured by neighbourhood preservation score (calculated for the first 150 genes). UMAP coordinates themselves were calculated using the whole transcriptome. For visualisation purposes, cell neighbourhood preservation scores lower than 0 were set equal to 0. Lower panel: UMAP representation coloured by celltype labels. Colours correspond to cell type labels and match colors in the upper panel.

We hypothesise that the difference in performance arises due to the nature of the L1-regularisation utilised by SCMER. In general, regularisations like Lasso tend to discourage redundancy among the selected features. However, this may not hold if the strength of the regularisation is chosen manually and lies above an appropriately selected range ^40^. We empirically support this hypothesis by assessing the redundancy within selected gene panels for both geneBasis and SCMER (**Supp. Fig. 2**). Specifically, when a low number of genes is selected, SCMER tends to select highly co-expressed genes.

Additionally, from the beginning of the search, geneBasis prioritises genes that allow delineation of cell types as well as scGeneFit, a method that specifically addresses the task of cell type separation (**Fig. 2B)**. Moreover, geneBasis robustly recovers genes that delineate cell types even if a preselected set of genes if provided (Methods, **Supp. Fig. 4**). Taken together, we suggest that geneBasis represents an efficient method to successfully resolve cell types and to preserve the overall manifold.

Another essential aspect of targeted gene panel design is to determine whether enough genes have been selected to capture variability in the dataset of interest. To this end, the cell neighbourhood preservation metric can be exploited to investigate whether specific transcriptional neighbourhoods are enriched for cells with low scores, suggesting that these neighbourhoods are not preserved well by the current selection. Across the benchmarked datasets, we illustrate this by focusing on the first 150 selected genes (**Fig. 2C, Supp. Fig. 3**). We observe that although there exist subtle differences between cell types, the majority of cell types (**Fig. 2C, upper panel**) and regions of the high-dimensional space (**Fig. 2C, middle panel**) were well covered by the selected panels, indicative of good performance (the only striking exception was in the melanoma dataset, in particular for cells that could not be assigned a clear label in the original analysis). Consistent with these results, we also observed high imputation accuracy for the expression of genes that were not among the selected set (**Supp. Fig. 3B,C**). More generally, we anticipate that a user may not necessarily want to select an exact number of genes, but rather may have an upper limit of genes that they could feasibly profile. In these cases, we advise running geneBasis until the limit is reached, before assessing when the cell neighbourhood preservation and gene prediction score distributions converge as a function of the number of genes (**Fig. 2, Supp. Fig. 3**). Finally, it is important to stress that the gene panel evaluation is independent of the selection method employed, and can be used to compare two or more panels generated by any approach.

### geneBasis accounts for batch effect and handles unbalanced cell type composition

Careful consideration and adjustment for batch effects is one of the most challenging aspects of scRNA-seq data analysis ^41^. In the context of gene selection, SCMER performs gene selection for each sample individually, and then retains features identified in all/most samples. While this is an efficient way to discard genes that show technical variability, such an approach might also discard genes with strong biological relevance that are only captured in a small fraction of samples e.g. cell type markers for cell types present in <50% of the batches. Importantly, such unbalanced cell type composition is not uncommon in practice.

To account for this, in cases where a batch is specified by a user (i.e. every cell is assigned to a batch), we construct k-NN graphs (‘true’ and ’selection’) within each batch, thereby assigning nearest neighbours only from the same batch and mitigating technical effects. Minkowski distances per genes are calculated across all cells from every batch thus resulting in a single scalar value for each gene. Importantly, if a certain cell type is present only in one batch, cells from this cell type will be consistently ‘mismapped’ if none of its marker genes are selected, and therefore the algorithm will select one of the marker genes.

To assess our approach, we utilized the first dataset described above, which consisted of four independent batches (samples) from the same stage of mouse development (**Supp. Fig. 5**). In each batch, a considerable fraction of cells were associated with blood lineage (11-27%, Methods). Next, we artificially removed cells in a batch-aware manner, thus keeping cells from the blood lineage in 1, 2 or 3 batches, as well as adding a positive control (retaining blood lineage in all the batches) and a negative control (removing blood lineage from all the batches). We applied geneBasis and SCMER to these different settings. geneBasis efficiently (among first 10 genes) selected one of the blood markers for each combination except for the negative control (**Supp. Fig. 5C**). SCMER identified strong markers in the positive control setting and when the blood lineage was retained in at least 2 of the 4 samples. However, it did not select any blood markers when the blood lineage was retained in only one sample (overall fraction of blood lineage cells in this combination was nearly 5%), even though the gene panel provided by SCMER was identified by the algorithm as of satisfactory quality (all selected genes for this case and KL divergence are listed in **Supplementary Table 3**).

### Computation complexity of geneBasis

With rapid increases in the size of scRNA-seq datasets, computational complexity of any algorithm is of increasing importance. Consequently, we sought to estimate the running time of geneBasis, to ensure its scalability. We established that three parameters are relevant for the computational complexity of the algorithm: **n** - number of cells, **D** - number of pre-filtered genes that undergo gene search, and **p** - number of genes we want to select. In practice, **p** will almost never exceed a few hundreds and **D** will never exceed 15000-20000, meaning that the limiting parameter is the number of cells. Accordingly, we estimated running time for a series of spleen scRNA-seq sub-datasets (Methods), whilst varying **n** from ∼5000 to ∼30000. We also varied **D** from 1000 to 10000, and **p** from 50 to 150. For the selected range, we observe a linear relationship between elapsed time and number of cells, and exponential increase in complexity as a function of **p** (**Fig. 3A**). As anticipated, change in **n** had a higher effect on the running time compared to change in **D** (**Fig. 3B**). Additionally, we compared computational complexity between geneBasis and SCMER, and established that across the tested range of **n, p** and **D** we outperform SCMER (**Fig. 3C**, for the details of the comparison see Materials and Methods). Note that this might not hold true for higher values of **p**, but since for all tested datasets selections with 150 genes appeared to be sufficient and complete (**Fig. 2**), we suggest that **p** <= 150 is a reasonable assumption. We further suggest that if a greater reduction in computational complexity is desired, appropriate downsampling strategies can be applied (Supplementary Note 1).

**Figure 3.**
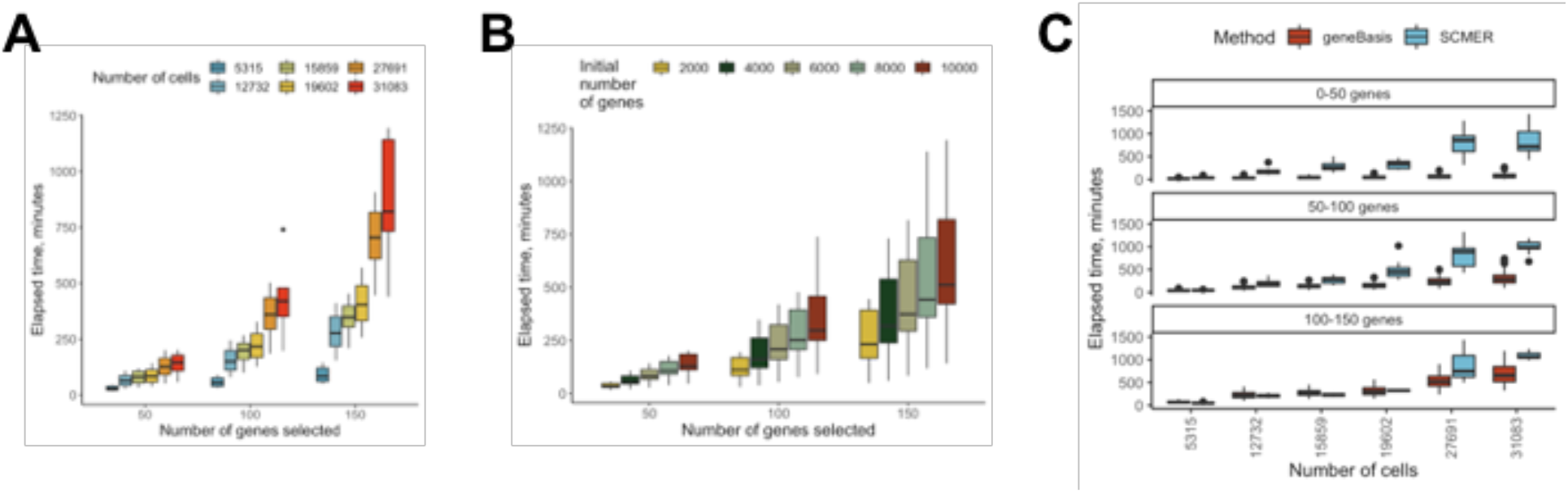
Estimation of computational complexity. (A) Distribution of elapsed time as a function of number of geneBasis selected genes (X-axis) and number of cells present in a scRNA-seq dataset (in colour). (B) Distribution of elapsed time as a function of number of geneBasis selected genes (X-axis) and initial number of genes (in colour). (C) Distribution of elapsed time as a function of number of selected genes (in facets) and number of cells (X-axis). Colours correspond to different methods.

### geneBasis resolves rare cell types and unannotated inter-celltype variability

Cellular identity, which is typically represented on the level of cell type, is a basic and essential unit of information that any targeted gene panel needs to recapitulate. In other words, a ‘biologically’ complete gene panel should delineate all cell types, including rare ones, for which statistical methods can be occasionally underpowered. However, it also should be able to capture intra-celltype variability in cases where discrete clustering has not been performed to the appropriate resolution, and identify genes that display gradual changes in expression across the high dimensional space, consistent with developmental trajectories. To address whether geneBasis satisfies the above criteria, we focused on three datasets: mouse embryogenesis, human spleen and human pancreas. All datasets contain annotated cell type labels and involve combinations of abundant and rare cell types. Moreover, each dataset contains cell types that are distinct and closely related, making them highly appropriate for the task (**Fig 4. B, H, J**).

**Figure 4.**
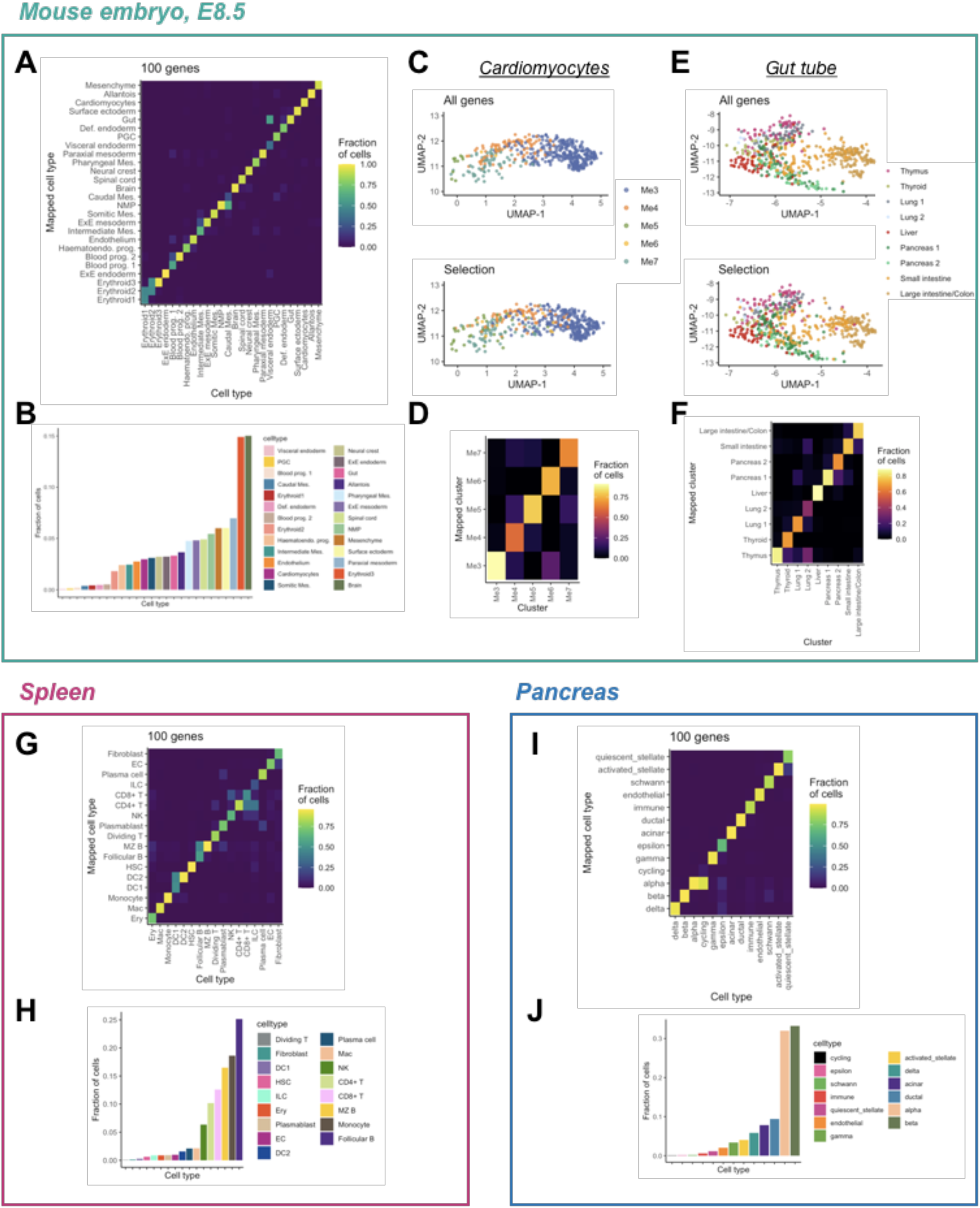
geneBasis delineates celltypes regardless of their abundance and resolves unannotated inter-celltype variability. Panels A-F correspond to mouse embryogenesis; G-H correspond to spleen, and I-J correspond to pancreas analysis. (A) Celltype confusion matrix for the first 100 selected genes (prog. = progenitors; mes. = mesoderm; Brain = Forebrain/Midbrain/Hindbrain). (B) Ordered bar plot representing relative celltype abundance. (C) Upper panel: UMAP representation of Cardiomyocytes, coloured by clusters from Tyser et al, when integration between datasets was performed using all genes. Lower panel: UMAP representation of Cardiomyocytes, coloured by mapped cardiac clusters, when integration between datasets was performed using selection from geneBasis. UMAP-coordinates themselves were calculated using the whole transcriptome. (D) Confusion matrix for cardiac clusters, where the mapping using all genes is used to assign true identity and mapping using the geneBasis selection is used to represent a mapped cluster. (E) Upper panel: UMAP representation of Gut tube, coloured by mapped organs (Nowotschin et al.), when integration between datasets was performed using all genes. Lower panel: UMAP representation of the Gut tube, coloured by mapped organs, when integration between datasets was performed using the selection from geneBasis. UMAP-coordinates themselves were calculated using the whole transcriptome. (F) Confusion matrix for the emerging organs present along the gut tube, where mapping using all genes are used to assign cells to a true cluster and the geneBasis selection is used to represent a mapped cluster. (G) Celltype confusion matrix generated using the first 100 selected genes. For cell types in spleen we use abbreviations, see Supplementary Table 2 for full annotations. (H) Ordered bar plot representing relative celltype abundance. (I) Celltype confusion matrix for the first 100 selected genes. (J) Ordered bar plot representing relative celltype abundance.

For the study of mouse embryogenesis, selecting only 20 genes accurately resolves most cell types (**Supp. Fig. 6B**), with 50-100 genes being sufficient to achieve nearly perfect matching (**Fig. 4A**). The minor exceptions comprise cell types that are transcriptionally similar in the original dataset, such as the visceral endoderm and the gut as well as the caudal mesoderm and the NMPs, where only few (if any) differentiating markers exist and cell types, and cell type classification is made using the degree of expression of commonly shared markers. Nevertheless, in all such cases geneBasis selects several genes that could in principle be used to differentiate the cell types (**Supp. Fig. 6D,E**).

Having determined that geneBasis identifies genes that characterise annotated cell types, we next asked whether it could find genes associated with more subtle biological signals present in the data. To this end we exploited recently published studies focused on myocardium development ^42^ and gut tube formation ^43^ (Methods). We first integrated (using all genes) cells annotated as cardiomyocytes from the scRNA-seq atlas for the whole embryo together with cells from ^42^ study that were isolated from the anterior cardiac region of mouse embryos, and assigned cluster identities to cardiomyocytes cells (Methods; **Fig. 4C**). As expected, all cells were assigned to the mesodermal cardiac lineage, with the majority being assigned to the most differentiated cardiomyocytes (Me3). Consistent with Tyser et al., 2021, the integrated data also reveal two trajectories leading to the most differentiated Me3 cluster: an FHF-like differentiation trajectory linking Me5, Me4 and Me3 and a SHF-like trajectory via Me7. Importantly, when using only the first 100 genes selected by geneBasis we were able to recapitulate this structure (**Fig. 4C, D**). Among the 100 genes, 13 were differentially expressed in cardiomyocytes (**Supp. Fig. 6F**), including markers for the FHF-trajectory (*Sfrp5*), strong markers for the differentiated Me3-state (*Myl2*) and contractile markers (*Ttn*) that exhibit gradual expression along the differentiation trajectory of the myocardium (**Supp. Fig 6G**). Similarly, when focusing on the gut tube, and using the refined annotation provided by ^43^, we observed that the set of genes chosen by geneBasis recovered the emerging organs arranged across the Anterior-Posterior axis (**Fig. 4E,F; Supp Fig 6H,I,K**).

Next, we explored the ability of geneBasis selection to select genes that captured heterogeneity amongst the populations of immune cells present in the adult human spleen. As expected, compared to mouse development, the ability to discriminate between the set of transcriptionally similar cell types present in the spleen required more genes (**Supp. Fig. 7B,C**). Nevertheless, selection of 100 genes allowed most cell types to be identified in a sensitive and specific manner (**Fig. 4G, Supp Fig 7D**). Similarly to mouse embryogenesis, the occasional mismapping of cells occurred between transcriptionally similar cell types, such as follicular and marginal zone B cells, as well as CD4+ and CD8+ T cells and innate lymphoid cells. Of note, the difficulty of using only the transcriptome to distinguish between different types of T cells is well known ^44^, and it has been suggested that a more precise annotation of T cells can be obtained by generating paired measurements of cellular transcriptomes and immunophenotypes ^45^.

Finally, we benchmarked geneBasis on a thoroughly annotated and integrated dataset of human pancreas, containing numerous rare cell types – 8 of the 13 annotated cell types account for less than 5% of the overall dataset. Nevertheless, and similarly to the previous examples, we quickly and accurately predict all cell types with the exception of rare cycling cells, which are occasionally conflated with highly abundant alpha cells (**Supp. Fig. 7F,G**).

### geneBasis captures signals associated with cell states

We next examined whether geneBasis could identify sources of biological variation that in principle can be recovered from scRNA-seq data but that are frequently overlooked because they do not contribute to cell identity per se. Examples of such genes include cell state markers associated with transient biological processes, such as cell cycle, DNA damage repair, oxidative stress. Depending on the biological context, this cell state information can be highly relevant for *in situ* profiling. For example, cell cycle genes are typically not included in gene panels for spatial transcriptomics experiments ^11,14,16^, partially due to high abundance of cyclin mRNAs and partially due to the notion that this signal is frequently deemed to be uninteresting. However, in the context of tumorigenesis, DNA damage, cell cycle and proliferation of malignant cells are the hallmarks of disease progression, and mapping this information spatially could be highly insightful.

To illustrate that geneBasis successfully and sufficiently recovers cell state genes indicative of ongoing biological processes, we analysed the set of genes recovered for the melanoma dataset (**Fig. 5A**), which consists of ∼4500 cells isolated from 19 patients, containing malignant, immune, stromal and endothelial cells. Within the first 100 genes selected, we identified 53 markers that were differentially expressed between the annotated cell types (Methods) and verified that these resolve all cell types, including the rare population of NK cells that are transcriptionally similar to highly abundant T cells (**Fig. 5A-C**).

**Figure 5.**
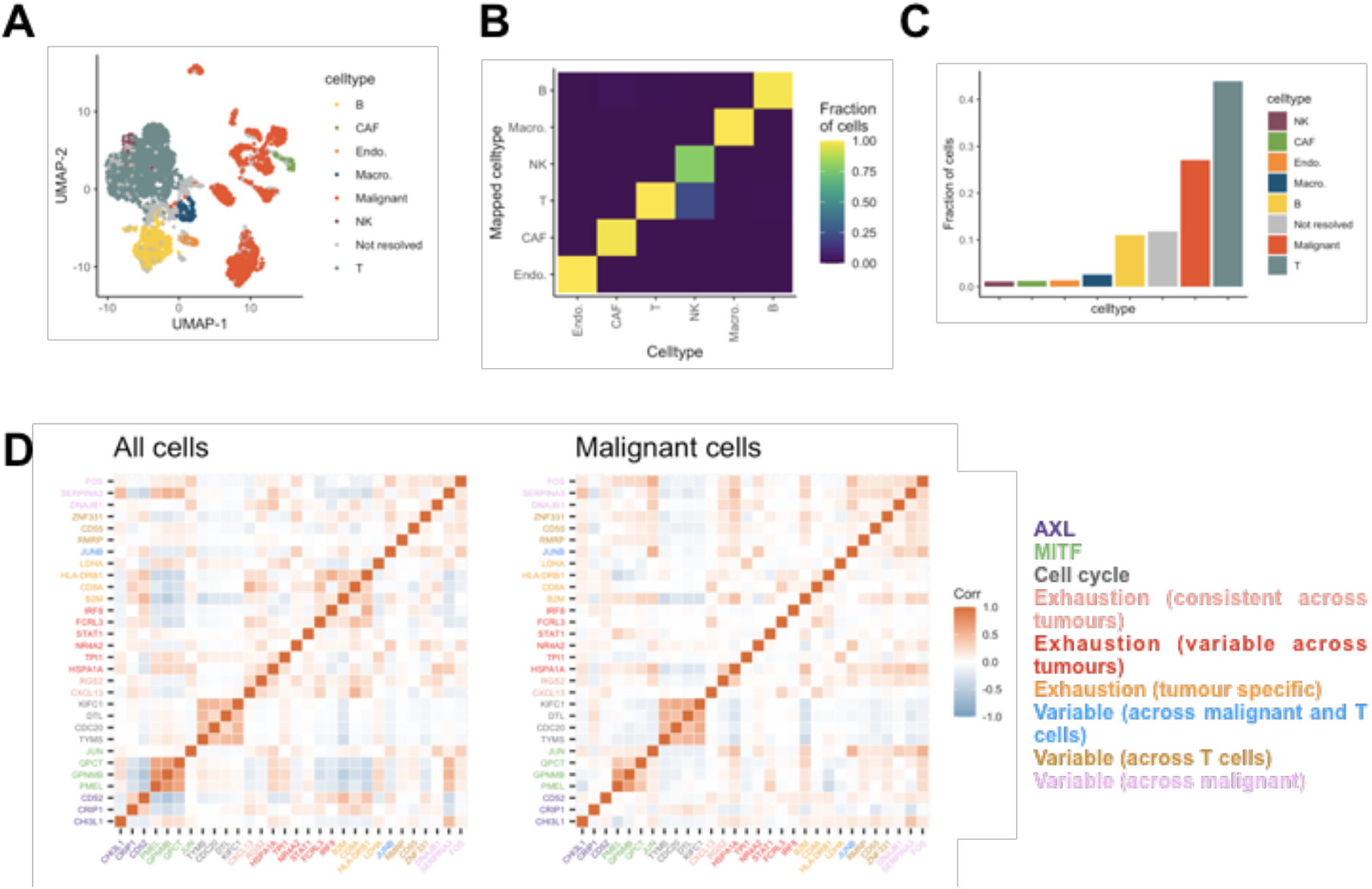
geneBasis identifies cell state genes relevant for heterogeneity across multiple celltypes in the context of cancer. (A) UMAP representation, coloured by malignancy status and annotated celltypes. UMAP-coordinates themselves were calculated using the whole transcriptome. (B) Celltype confusion matrix for non-malignant cells. (C) Ordered bar plot for celltype abundance. (D) Co-expression (within all cells, left panel; within malignant cells, right panel) for selected genes associated with various transcriptional programs identified in the original publication. Colours correspond to detected programs, corresponding legend is on the right.

The majority of the remaining genes were globally differentially expressed between malignant and non-malignant cells (8 and 26 genes were down- and up-regulated respectively in malignant cells). In the original study, a substantial degree of transcriptional heterogeneity was characterised across individual cells, both malignant and healthy, mostly associated with cell cycle, variation in MITF and AXL levels, and activation of the exhaustion program in T cells. Importantly, among the up-regulated genes we identified markers of transcriptional programs described in the original study such as tumour progression and cell exhaustion.

Finally, 13 genes were neither identified as cell type markers nor significantly up or down regulated in malignant cells. Among those we found *JunB, B2M* and *Narf2* – markers of inflammation, cell exhaustion and melanoma oncogenic programs taking place in malignant and T cells. Overall, we select cell state markers for all cell states characterised in the original publication programs including MITF and AXL (**Fig. 5D**). We observed a low degree of co-expression among these genes, with the exception of cell cycle genes and genes associated with the MITF program. Low co-expression is consistent both across all cells and within malignant cells, suggesting that these cell state genes are indicative of distinct transcriptional programs.

## Discussion

For the last decade, scRNA-seq technology has transformed our ability to explore molecular heterogeneity in a variety of biological systems. More recent technological advances such as single-cell multi-omics assays, CRISPR screens and spatial transcriptomics go beyond measuring only the transcriptome, thus facilitating a more complete understanding of the features that underpin cellular function. In many of these cases, particularly for a large number of spatial transcriptomics assays, selecting the set of genes to probe is an important parameter, which in turn necessitates the emergence of appropriate computational tools.

In this study, we introduce geneBasis, which uses existing scRNA-seq data to select genes in a cluster-free and highly flexible manner. We have shown that geneBasis outperforms existing methods, both in terms of computational speed and in identifying relevant sets of genes, and that geneBasis selects genes that characterise both local and global axes of variation that can be recovered from a k-NN graph representation of transcriptional similarities. geneBasis also allows user knowledge to be directly incorporated by selecting, *a priori*, a set of genes of particular biological relevance, which are then augmented by the algorithm.

Although we have addressed several important challenges, our approach has limitations that need to be considered, especially when designing libraries for FISH-based experiments. Firstly, scRNA-seq and FISH-based technologies use different approaches for assessing the number of mRNA molecules that are associated with a given gene, which in turn creates discrepancies in capture efficiency between the two technologies ^46^. Practically, it is observed that FISH-based technologies capture more mRNAs per cell due to a high sequence specificity in probe design. Consequently, genes selected using scRNA-seq will likely have higher detection rates in FISH-based experiments. While this is beneficial for detection of cell types that rely on a low number of selected markers, it can be a problem since FISH-based experiments are experimentally limited by the total number of individual molecules that can be detected in any given cell. Accordingly, it is advised to limit the number of ubiquitously or highly expressed genes. By design, geneBasis does not perform initial pre-filtering of housekeeping genes (since some could be highly relevant, such as markers of the cell cycle) nor it does discard them from the selected gene panels. Therefore, we suggest that users filter out ubiquitously expressed genes either in the initial submission or post hoc.

Another important aspect to be accounted for in the selection of gene panels is redundancy (i.e., the presence of multiple genes that are co-expressed and that mark the same cell type(s)). In practice, although geneBasis is not explicitly designed to completely eliminate redundancy in the selection, we typically observed only a small number of redundant genes. In general, this is a desirable property, since it maximises the number of distinct biological processes that can be examined. However, for cell types of specific interest, it may be useful to add a small number of redundant genes to the gene panel to avoid risks associated with technical probe failure during the experiment.

Finally, since geneBasis does not rely upon clustering of the scRNA-seq data, it does not directly capture ‘positive’ cell type markers. As an example, when examining the ability to detect primordial germ cells (PGCs), a rare population comprising ∼0.1% of cells in the mouse embryo study, we showed above that selecting 100 genes successfully resolves all cell types including PGCs. However, manual annotation of the selected genes revealed that we do not select unique positive PGC markers in the selected set. Instead, we select two semi-specific PGC markers, *Ifitm1* and *Phlda2*, which are shared with other cell types such as somitic and extra-embryonic mesoderm, extra-embryonic endoderm and allantois. The ability to discriminate PGCs from the other related cell types is possible due to the inclusion of genes that mark these related cell types as opposed to the inclusion of specific PGC marker genes (of note, amongst the top 200 genes we include *Dppa3* - a highly specific and sensitive PGC marker). Consequently, we suggest that in cases where sensitive and specific identification of cell types, particularly rare ones, is a high priority, that appropriate marker selection methods are applied alongside geneBasis, thus ensuring the inclusion of specific cell type markers.

## Methods

### Detailed overview of the geneBasis algorithm

Below we describe the workflow of the algorithm, characterise optional parameters and justify the default settings.

#### Initial selection of genes

We perform initial pre-filtering of genes by using the function `scran::modelGeneVar`, `var.threshold = 0`. If the number of variable genes exceeds `n = 10000`, we use the first 10000 variable genes as default. Additionally, we filter out mitochondrial genes. If a user wants to perform more selective filtering, this can be done manually (by using scRNA-seq entries for only pre-filtered genes) or by tuning `var.threshold` and `n`.

#### k-NN graph representations for ‘true’ and ‘selection’ graphs

- We use log-normalised counts for scRNA-seq data as an input for the algorithm.
- We construct kNN-graphs, for both true and selection graphs, by using the function `BiocNeighbors::findKNN,` and represent each cell’s neighbourhood using the first 5 neighbours. As a default, we use `n.neigh=5` as an appropriate compromise between mitigating potential noise in scRNA-seq data together with the potential existence of very rare cell types, where `n.neigh > 5` can hide true biological variability. The same default is used to compute the cell neighbourhood preservation and gene prediction scores. This variable can be tuned by the user
- For the ‘true’ graph, we first compute Principal Components by using the function `irlba::prcomp_irlba`, `n = 50`, and apply `BiocNeighbors::findKNN` on the first 50 PCs.
- For the ‘selection’ graphs, the PCA step is optional and by default is not performed. In practice we observed that omitting the PCA step for the construction of ‘selection’ graphs did not lead to different results when selections are generated with fewer than 250 genes (**Supp. Fig. 9**).

#### Identification of the first gene

To select the first gene, we generate a random gene count matrix thus mimicking a random transcriptional relationship between cells. Subsequently, we use this matrix to compute a random ‘selection’ graph. Subsequently, to calculate the per gene Minkowski distance in the random ‘selection’ graph, we compute the distance between the actual log-normalized expression values and the average of the ‘true’ log-normalized expression values across neighbours in the random ‘selection’ graph. To select the first gene we choose the gene with the largest difference between Minkowski distance in the random ‘selection’ graph and the ‘true’ graph. We repeat this procedure five times and select the most frequently occurring gene. In the unlikely scenario of ties (5 different genes are calculated from 5 random graphs), we select the gene from the first random graph.

#### Adding a new gene to the current gene panel

To add the next gene to the current gene panel (selection), we compare the graph constructed using the entire transcriptome (‘true’ k-NN graph, computed once) with a graph constructed using the current ‘selection’ (‘selection’ graph, recomputed for every updated selection). For each gene in the ‘true’ graph we compute, across cells, the Minkowski distance between a gene’s expression in a given cell and its average expression in that cell’s k nearest neighbours. Similarly, we compute the Minkowski distance for each gene using the ‘selection’ graph. To add the next gene to the current gene panel, for each gene we calculate the difference in stance between ‘selection’ and ‘true’ graph, and add the gene with the largest difference.

#### Batch correction

In cases where a batch is specified by a user (i.e. every cell is assigned to a distinct batch), we construct k-NN graphs (‘true’ and ‘selections’) per batch thus identifying nearest neighbours only from the same batch. To calculate a per gene Minkowski distance between cells and their neighbours, we use cells from all batches. Note that the same accounting of the batch is performed to compute cell neighbourhood preservation and gene prediction scores.

#### Default order for Minkowski distance

The Minkowski distance of order *p* (*p* is an integer) between two vectors *X* = (*x*_*1*_, *x*_*2*_, *…, x*_*n*_) and *Y* = (*y*_*1*_, *y*_*2*_, *…, y*_*n*_) ∈ *R*^*n*^ is defined as:

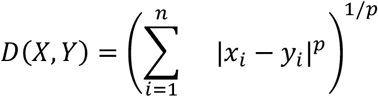

The choice of *p* provides a balance between how much the computed distance weighs the number of cells that are currently ‘mismapped’ (i.e. the assigned neighbours in the selection graph show different expression values from what would be expected from the ‘true’ graph) against the magnitude of this ‘mismapping’ (i.e. how much different). For example, when the Minkowski distance is calculated between vectors *X* and *Y* and *p* → ∞, then *D*(*X, Y*) → (|*X*_*i*_ *− Y*_*i*_ |) thus prioritising the magnitude of mismapping. By contrast, if *p* = 0 then the Minkowski distance emphasises the number of mismapped cells regardless of the magnitude.

To select an appropriate *p*, we applied geneBasis for the range `p = 1,2,3,4,5` (**Supp. Fig. 8**).Overall, we observe that different orders of Minkowski distance return robust selections that do not substantially affect either cell neighbourhood preservation or gene prediction scores (**Supp. Fig. 8A,B**). We next sought to determine the degree of expression resolution that can be achieved for a given range of *p*. In other words, we assessed, for each value of *p* and varying number of selected genes, what is the most rarely expressed gene (measured as a fraction of cells in which non-zero counts are observed) can be achieved. We observe that for all benchmarked datasets *p* > 2 gives highly similar results and using *p* > 2 allows the selection of relevant genes that are expressed in only ∼0.1% of cells (**Supp. Fig. 8C**). Applying geneBasis with `p = 1,2` shows less flexibility and does not select genes beyond a certain threshold of expression. As default, we thus select `p = 3` as a good tradeoff between the number of mismapped cells and the size of the discrepancy.

### Gene prediction score (‘gene score’)

To calculate the gene score for the selected library, we exploit the ‘selection’ and ‘true’ graphs. For each gene, we compute the Spearman correlation (across cells) between its log-normalised expression value in each cell and its average expression across (each cell’s) first 5 nearest neighbours. We perform this calculation separately for the ‘selection’ and ‘true’ graphs and compute the ratio [correlation in ‘selection’]/[correlation in ‘true’]. We discard genes with correlation in the ‘true’ graph below 0.25 (i.e. we omit genes unstructured expression across the manifold).

### Cell neighbourhood preservation score (‘cell score’)

To calculate the cell score, we exploit the ‘True k-NN graph’. In the ‘True k-NN graph’ for each cell we compare the vector of distances to the assigned neighbours from the ‘true graph’ and assigned neighbours from the ‘selection graph’ (**Supp. Fig. 2**). To normalize vectors and provide an interpretable metric, for each cell we z-transform the distribution of distances to all other cells and multiply the vector by -1, thus higher distances reflect more transcriptionally similar cells. For each cell, the ‘cell score’ is calculated as the ratio of the median of the scaled distances to the first K neighbours from the ‘selection’ graph to the median across of the scaled distances to the first K neighbours from the ‘true’ graph (we use K = 5). The upper limit for the score of 1 arises if a cell’s neighbourhood perfectly overlaps between the ‘true’ and the ‘selection’ graph, while a score of 0 suggests that a cell’s neighbours from the ‘selection’ graph are distributed randomly across the manifold.

### Cell type mapping

To compute confusion matrices for cell type labelling using the selection, we use log-normalised counts of the selected genes, perform PCA and use the first 50 components to build a k-NN graph. If batch is not provided, PCA is performed using the `prcomp_irlba` function from the `irlba` package. If batch is provided, we first perform PCA using the `multiBatchPCA` function from the `batchelor` package, and then perform MNN-correction using the `reducedMNN` function from the `batchelor` package. For each cell we assign the mapped cell type as the most frequent cell type label across the first 5 neighbours (in case of the ties, we assign the cell type label of the closest cell from tied cell types).

### Benchmarking

#### Datasets

- <u>Mouse embryogenesis.</u> The dataset was downloaded using the `MouseGastrulationData` package in R, which contains data generated for ^32^. Further we selected samples for stage E8.5 and discarded cells annotated as doublets or stripped. The final dataset consists of 4 samples (i.e. batches), and we use the field “sample” as a batch identification for gene selection methods. For cell type mappings we discard extremely rare cell types that are present only at earlier developmental stages: Paraxial Mesoderm, Notochord and Rostral Neuroectoderm (< 10 cells in the E8.5 dataset).
- <u>Spleen.</u> scRNA-seq for spleen is downloaded from the HubMap portal ^47^ (https://portal.hubmapconsortium.org). The HuBMAP dataset IDs are as follows: HBM984.GRBB.858 HBM472.NTNN.543 HBM556.QMSM.776 HBM336.FWTN.636 HBM252.HMBK.543 HBM749.WHLC.649. Raw counts were SCTtransformed and integrated using Seurat RPCA workflow. Annotation of cell types was performed using a manually curated list of previously characterised cell type markers (Supplementary Table 2 contains the list of spleen markers and full annotations). Cells that could not be confidently assigned to a single cell type based on these markers were denoted as “Unknown.” The dataset consists of samples from 6 donors (i.e. batches), and we use the field “donor” as a batch identifier for gene selection methods.
- <u>Pancreas.</u> Annotated scRNA-seq datasets for the pancreas were obtained from the Azimuth portal ^45,48^. The integrated dataset contains data from 6 individual studies ^33– 38^, with each individual study also being composed of several batches or samples. We use a combination of study and batch within study as a meta-batch variable (20 in total).
- <u>Melanoma.</u> The melanoma dataset described by ^39^ was downloaded from ncbi.nlm.nih.gov with accession number GSE72056. The dataset consists of samples from 19 donors at different stages of metastasis. Considering the highly variable and overall low number of cells per each donor (4645 cells across 19 donors), and, more importantly, substantial transcriptional variability across donors, we decided to perform gene selections for this study without performing batch correction. For cell type mappings, we only use annotated non-malignant cells.

#### Pre-processing

To obtain log-normalised counts we first calculated size factors using `scran::quickCluster` to cluster cells and subsequently `scran::computeSumFactors` to compute size factors. Log-normalisation was then performed using `batchelor::multiBatchNorm`.

### Benchmarked methods

- SCMER and scGene-Fit require initial pre-selection of genes. We used HVG-based gene screening with defaults from SCMER and manually increased the search range for spleen due to the observation that the defaults in SCMER selected an insufficient number of genes that poorly represented the overall manifold. Other parameters of the algorithms were left as default.
- For SCMER, we used the same batch identifications as for geneBasis. To get a range of gene selections for SCMER, we generated a coarse grid for regularisation strength (the parameter that directly affects number of genes selected) and applied the method on the generated grid.
- In order to perform a faithful comparison between scGene-Fit and geneBasis, for scGene-Fit we discarded the Unknown cell type for the spleen data and Paraxial Mesoderm, Notochord and Rostral Neuroectoderm cell types for the Mouse embryo data. For all 4 datasets, we used the option “centers”.

### Estimation of computation complexity for geneBasis and SCMER

geneBasis and SCMER use distinct computational strategies that required us to approach the estimation of computational complexity separately for each method. To vary the number of cells, we used the spleen dataset containing samples from 6 donors (henceforth referred as samples). We applied both methods to subsets of the spleen dataset by varying which samples we included in the subsets. Below we explain how we then assessed computational complexity for both methods.

- *<u>geneBasis</u>*. To vary the initial number of genes we performed highly variable gene selection and selected the top 2000, 3000, 4000, 5000, 6000, 7000, 8000, 9000 or 10000 highly variable genes. We then ran the algorithm to select the first 150 genes, and accordingly for every added gene, recorded the elapsed time.
- *<u>SCMER</u>*. SCMER requires the regularization strength λ to be inputted. Subsequently, given this parameter, a selected set of genes are returned. Alternatively, it can take a number of genes to be selected as an input, in which case it will run SCMER on the coarse grid of various values for λ, select an optimal λ to achieve the desired number of genes, and return the selection. By design, the latter takes longer to run since the algorithm has to first perform a grid search. Accordingly, to achieve a more faithful comparison, we selected the appropriate grid for λ ourselves, and for each value of the grid, we recorded the selected number of genes and elapsed time. The initial preselection of genes is not part of SCMER, and we varied the number of initial genes by selecting highly variable genes with `scanpy.pp.highly_variable_genes` with different values for `min_disp` (0, 0.25, 0.5).

### Generation of semi-random initial selections (for Supplementary Figure 4)

To perform semi-random initial selections, for each dataset, we randomly chose half of the cell types and for each cell type we randomly selected a gene that was upregulated in the corresponding cell type. A union of these genes was used as initial selections.

### Integration with lineage-specific datasets for mouse embryo development

To integrate scRNA-seq data of the whole mouse embryo ^32^ together with the lineage specific dataset for cardiac development ^42^, we concatenated log-normalised counts for cells from ^32^ annotated as ‘Cardiomyocytes’ and cells from ^42^. To integrate scRNA-seq data of the whole mouse embryo ^32^ together with lineage specific dataset for endodermal development ^43^, we concatenated log-normalised counts for cells from ^32^ annotated as ‘Gut’ and cells collected at E8.75 from ^43^.

For both integrations, we performed cosine normalisation of log-normalised counts using `batchelor::cosineNorm`, and performed PCA using `batchelor::multiBatchNorm`, 20 PCs (using combination of + dataset + corresponding to the dataset batch entity as meta-batch). Mapping was then calculated using `BiocNeighbors::findKNN`. For each cell, we assigned the most common cluster across the first 5 neighbours.

## Acknowledgements & Funding

We thank Emma Dann and Mike Morgan for useful discussions on the package development. This work is supported by the National Institutes of Health: RS and JM acknowledge 1OT2OD026673-01 which supports AM; RS acknowledge RM1HG011014-02; TS acknowledges K99HG011489-01; MA, HN, CW and TB acknowledge U54AI142766. SG acknowledges Royal Society Newton International Fellowship (NIF\R1\181950). MA and TB acknowledge Helmsley Charitable Trust 2004-03813. JM acknowledges core funding from EMBL and core support from Cancer Research UK (C9545/A29580). In the past three years, RS has worked as a consultant for Bristol-Myers Squibb, Regeneron, and Kallyope, and served as an SAB member for ImmunAI, Resolve Biosciences, Nanostring and the NYC Pandemic Response Lab.

## Author contributions

AM conceived the method, developed the algorithm, performed the analysis and wrote the R package. MB, CW, HN, TB & MA generated scRNA-seq data for the adult spleen. JJ preprocesed and annotated spleen scRNA-seq data. AB integrated pancreas data from the original publications. RS, JJ, AB, SG and TS provided critical feedback and helped shape the methodology, analysis and the package. AM and JM wrote the manuscript with the input from RS, JJ, AB, SG and TS. RS and JM oversaw the project.

