## Supplementary note 1 for "geneBasis: an iterative approach for unsupervised selection of targeted gene panels from scRNA-seq"

**geneBasis is robust to downsampling as a strategy to reduce computational complexity.**

Computational complexity is an important aspect of all successful algorithms that are anticipated to be utilized routinely. In the case of geneBasis, we suggest that if a reduction in computational complexity is necessary, downsampling of cells strategies can be applied.

To support this notion and to demonstrate the robustness of geneBasis to downsampling, we applied it to spleen data from individual samples (**Supp. Notes, Fig. 1**). The cell type composition across samples was fairly balanced, however, a certain degree of variation was observed for individual cell types, including the complete absence of 'Fibroblasts' in one of the samples (**Supp. Notes, Fig. 1A**). Importantly, although using a much smaller number of cells, we observe that selections that were generated from individual samples resolve cell types (**Supp. Notes, Fig. 1B**) and preserve neighbourhoods (**Supp. Notes, Fig. 1C,D**) as well as the selection generated using all samples.

We suggest that with a cautious downsampling strategy, preferably ensuring inclusion of all annotated cell types if annotations exist, the computation complexity of geneBasis can be reduced significantly. Finally, if only one/few cell types are of particular interest, geneBasis can be performed directly on those cell types which will greatly decrease computational complexity.

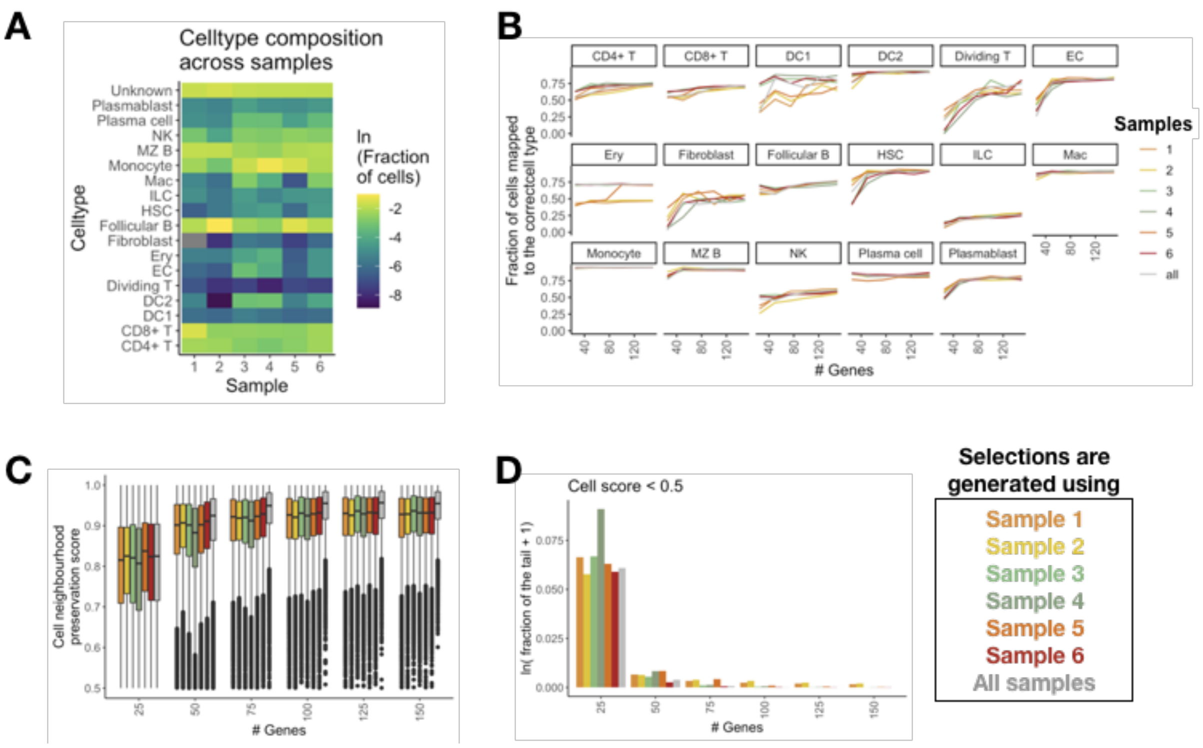


**Supplementary Notes Figure1. geneBasis provides robust and highly comparable and similar selections for downsampled datasets.** Gene selection was performed using each spleen sample (i.e. batch) separately as well as using the combined dataset.

(A) Celltype composition across samples (donors) indicates a certain degree of variation for individual celltypes.

(B) Per celltype, X corresponds to the number of the genes in the panel and Y corresponds to the fraction of correctly mapped cells. Bright colors from yellow to red correspond to the selections generated using individual samples and gray corresponds to the selection using all cells.

(C) Cell neighborhood preservation score distribution across selections and numbers of genes in the panel.

(D) Weights of the tails for cell neighborhood preservation score distributions (fraction of cells with neighborhood preservation score < 0.5) across selections and numbers of genes in the panel.
