## Supplementary Figures for "geneBasis: an iterative approach for unsupervised selection of targeted gene panels from scRNA-seq"

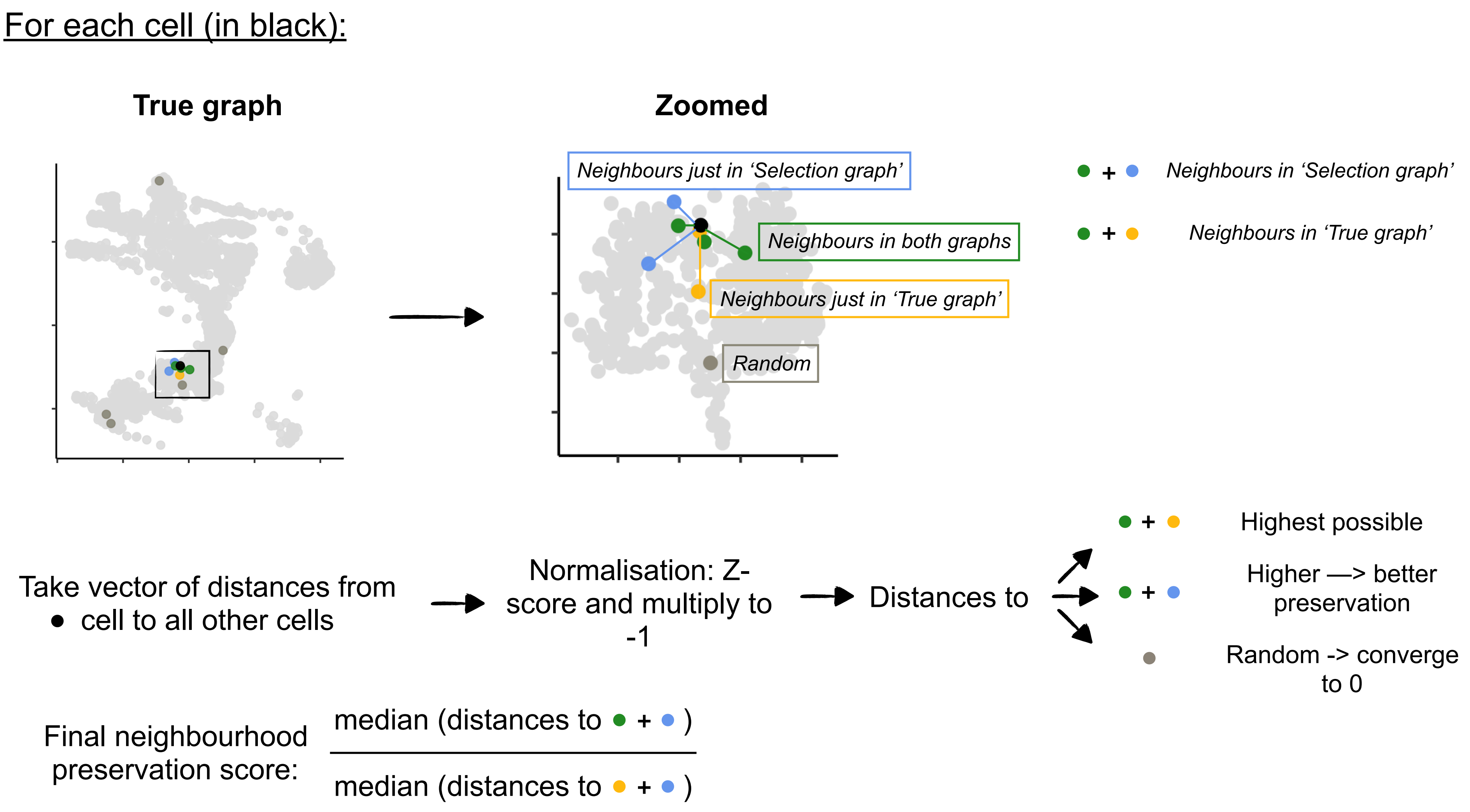


**Supplementary Figure 1. Schematic visualisation for cell neighbourhood preservation score workflow.** For each cell (in black) we detect neighbours from the True graph and from the Selection graph, and use the ratio of normalised distances as the final score.

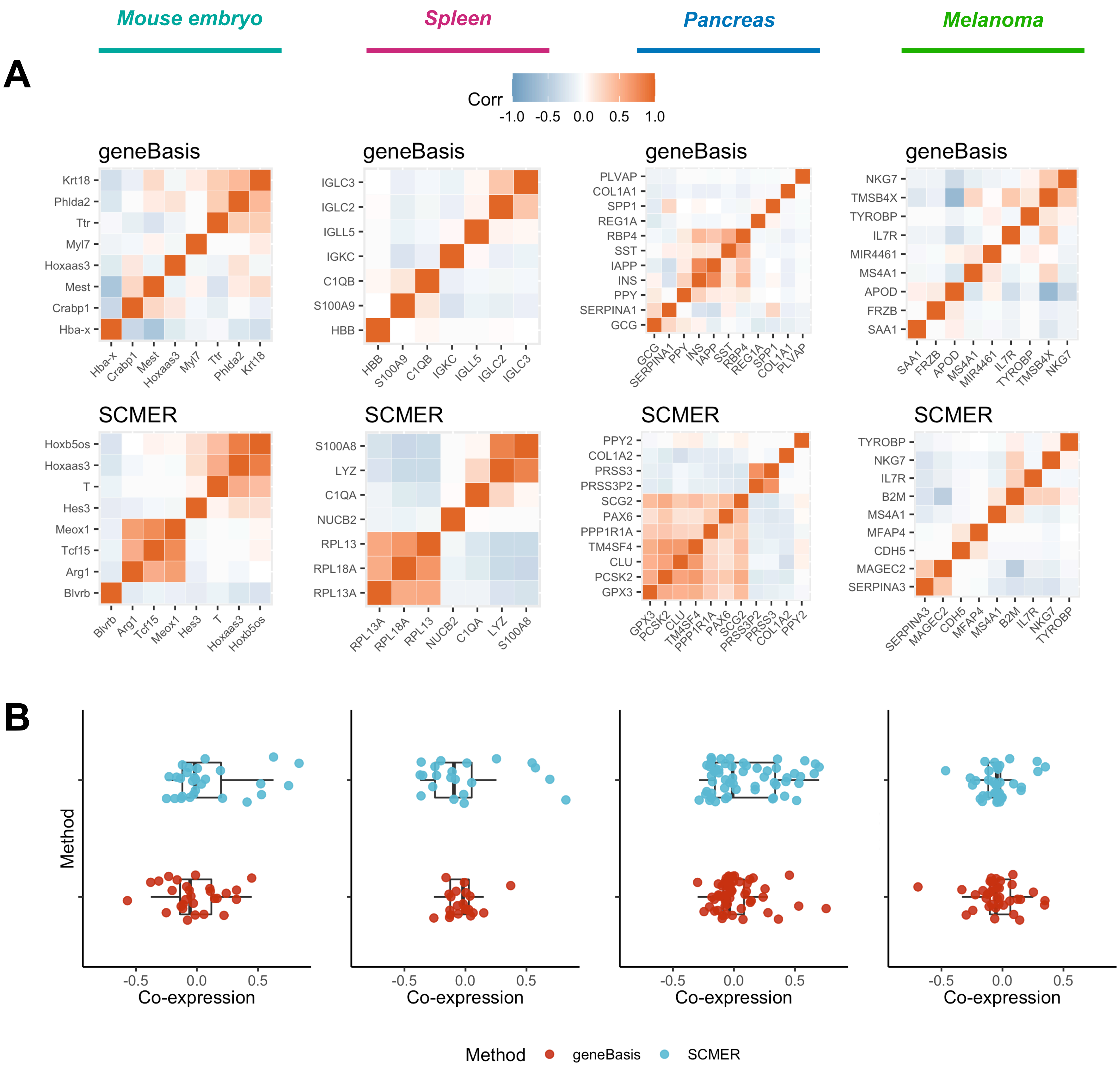


**Supplementary Figure 2. SCMER introduces redundancy in the selections when selecting a small number of genes.**

(A) Heatmaps showing the co-expression (Spearman correlation) between genes selected by geneBasis and SCMER. The overall degree of co-expression is lower for geneBasis compared to SCMER, and this is consistent for all benchmarked datasets.

(B) Boxplots represent pairwise correlation in log-normalized expression values (Y-axis) for the first genes selected for geneBasis (red) and SCMER (light blue).

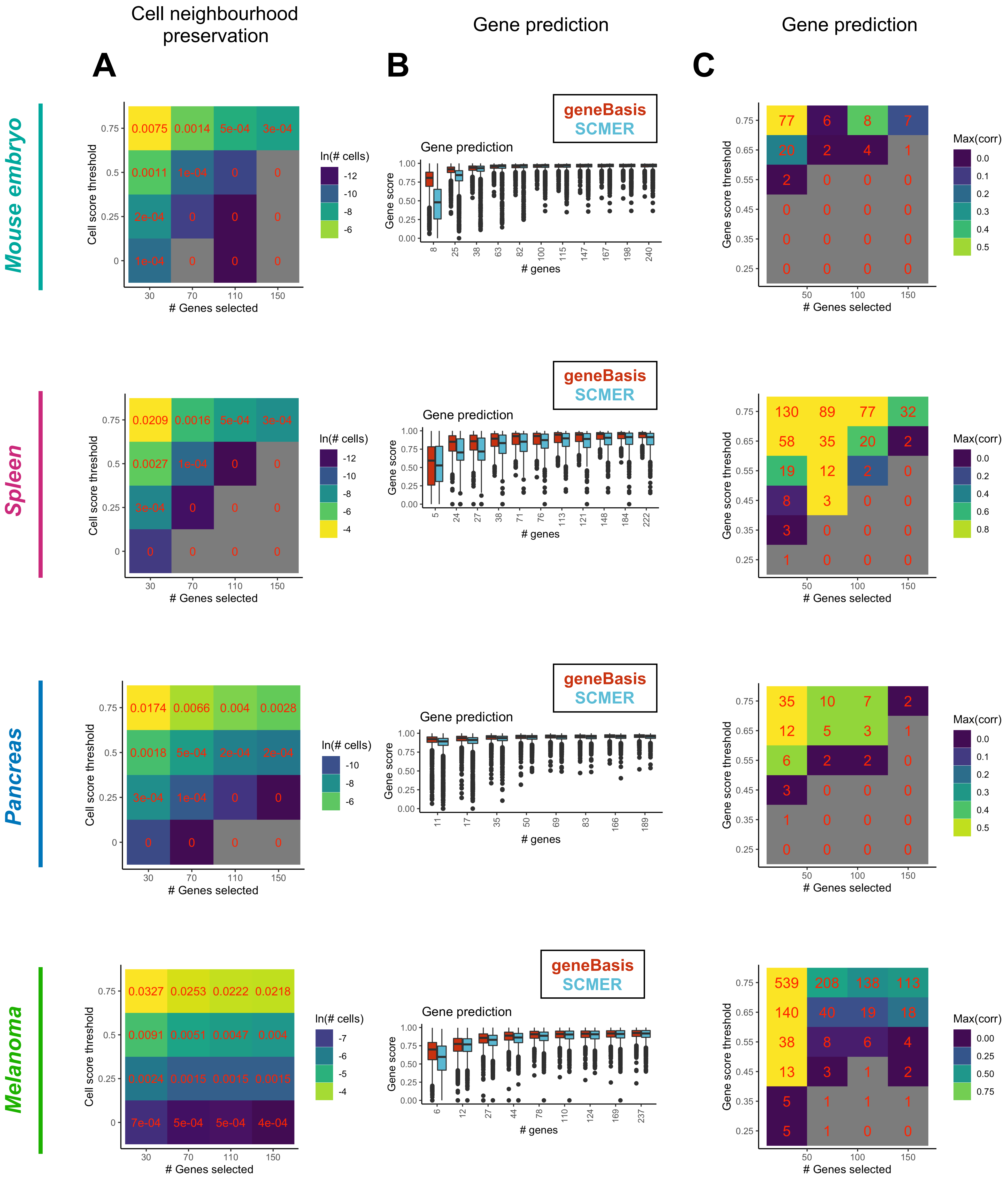

Cell neighbourhood preservation
Gene prediction
Gene prediction
A
B
C
Mouse embryo
Spleen
Pancreas
Melanoma
Cell score threshold
### Genes selected
ln(# cells)
geneBasis
SCMER
Gene score
### genes
Gene score threshold
Max(corr)

**Supplementary Figure 3. Systematic assessment of the ‘completeness’ of the generated gene panels. We provide several semi-orthogonal metrics to evaluate how complete the current panel is. Analysis is performed for each dataset (in rows).**

(A) Heatmap represents the fraction of cells that have low neighbourhood preservation score - as a function of number of genes (X-axis) and threshold for neighbourhood preservation score (Y-axis). Colours are log-scaled to facilitate visualization. Numbers inside correspond to actual values.

(B) Distribution of gene prediction score as a function of the gene panel size. Colours correspond to different methods.

(C) Heatmap represents statistics regarding gene prediction score - as a function of number of genes (X-axis) and threshold for gene prediction score (Y-axis). Numbers inside correspond to the number of genes that show a gene prediction score lower than the corresponding threshold. Colours correspond to maximum across correlations for all pairwise comparisons between genes with low gene prediction scores. Colours are log-scaled to facilitate visualisation.

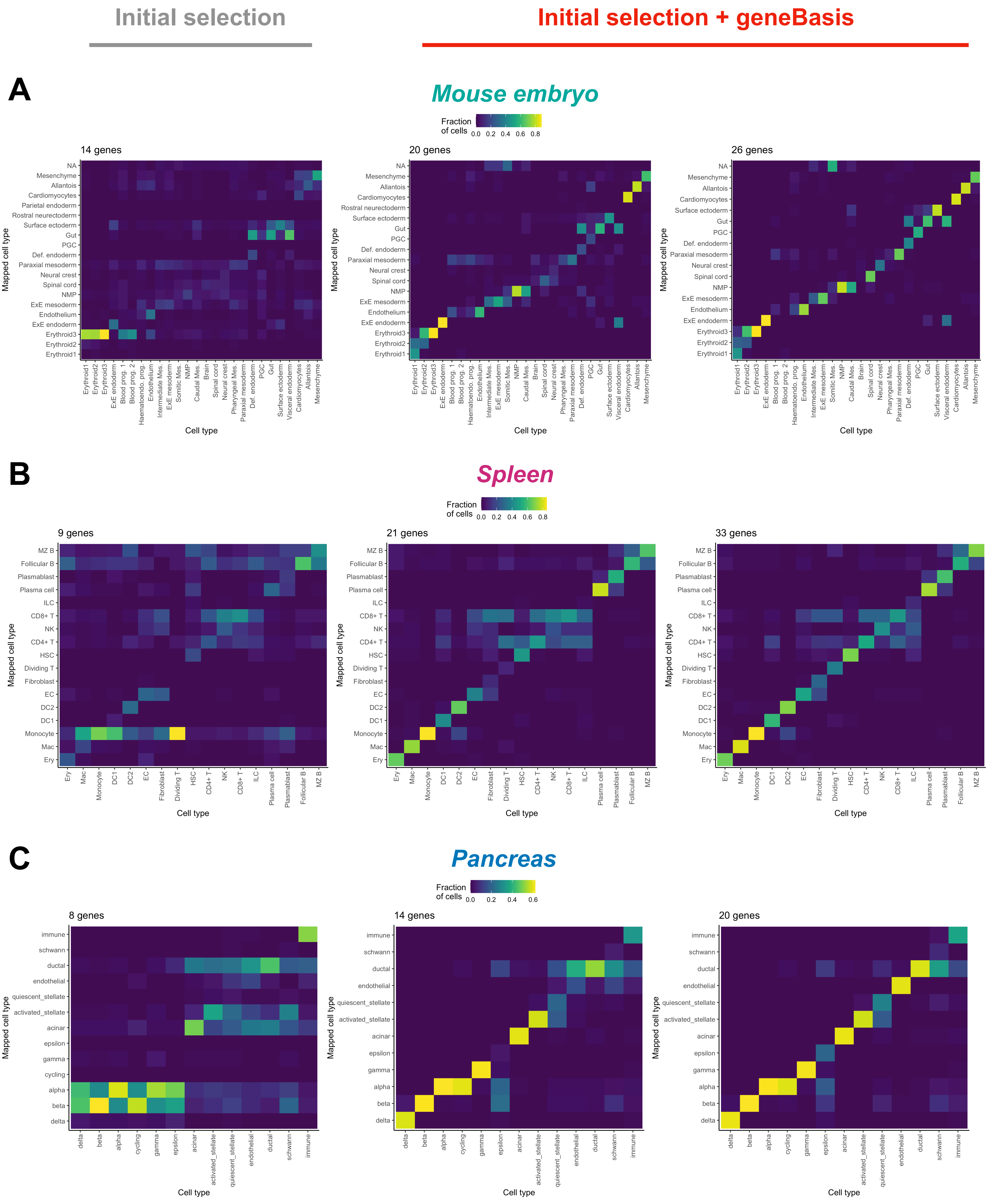
Initial selection
Initial selection + geneBasis
A
Mouse embryo
Fraction of cells
0.0 0.2 0.4 0.6 0.8
14 genes
20 genes
26 genes
Mapped cell type
Cell type
NA
Mesenchyme
Allantois
Cardiomyocytes
Parietal endoderm
Rostral neurectoderm
Surface ectoderm
Gut
PGC
Def. endoderm
Paraxial mesoderm
Neural crest
Spinal cord
NMP
ExE mesoderm
Endothelium
ExE endoderm
Erythroid3
Erythroid2
Erythroid1
Blood prog. 1
Blood prog. 2
Haematoendo. prog.
Intermediate Mes.
Somitic Mes.
Caudal Mes.
Brain
Pharyngeal Mes.
Visceral endoderm
B
Spleen
Fraction of cells
0.0 0.2 0.4 0.6 0.8
9 genes
21 genes
33 genes
Mapped cell type
Cell type
MZ B
Follicular B
Plasmablast
Plasma cell
ILC
CD8+ T
NK
CD4+ T
HSC
Dividing T
Fibroblast
EC
DC2
DC1
Monocyte
Mac
Ery
C
Pancreas
Fraction of cells
0.0 0.2 0.4 0.6
8 genes
14 genes
20 genes
Mapped cell type
Cell type
immune
schwann
ductal
endothelial
quiescent_stellate
activated_stellate
acinar
epsilon
gamma
cycling
alpha
beta
delta

**Supplementary Figure 4. geneBasis is robust to initial selections and quickly finds missing sources of variation.**

(A) Mouse embryo. Celltype confusion matrices for the initial semi-random selection (left), for the updated selection with 6 additional genes (middle) and updated selection with 12 additional genes (right).

(B) Spleen. Celltype confusion matrices for the initial semi-random selection (left), for the updated selection with 12 additional genes (middle) and updated selection with 24 additional genes (right).

(C) Pancreas. Celltype confusion matrices for the initial semi-random selection (left), for the updated selection with 6 additional genes (middle) and updated selection with 12 additional genes (right).

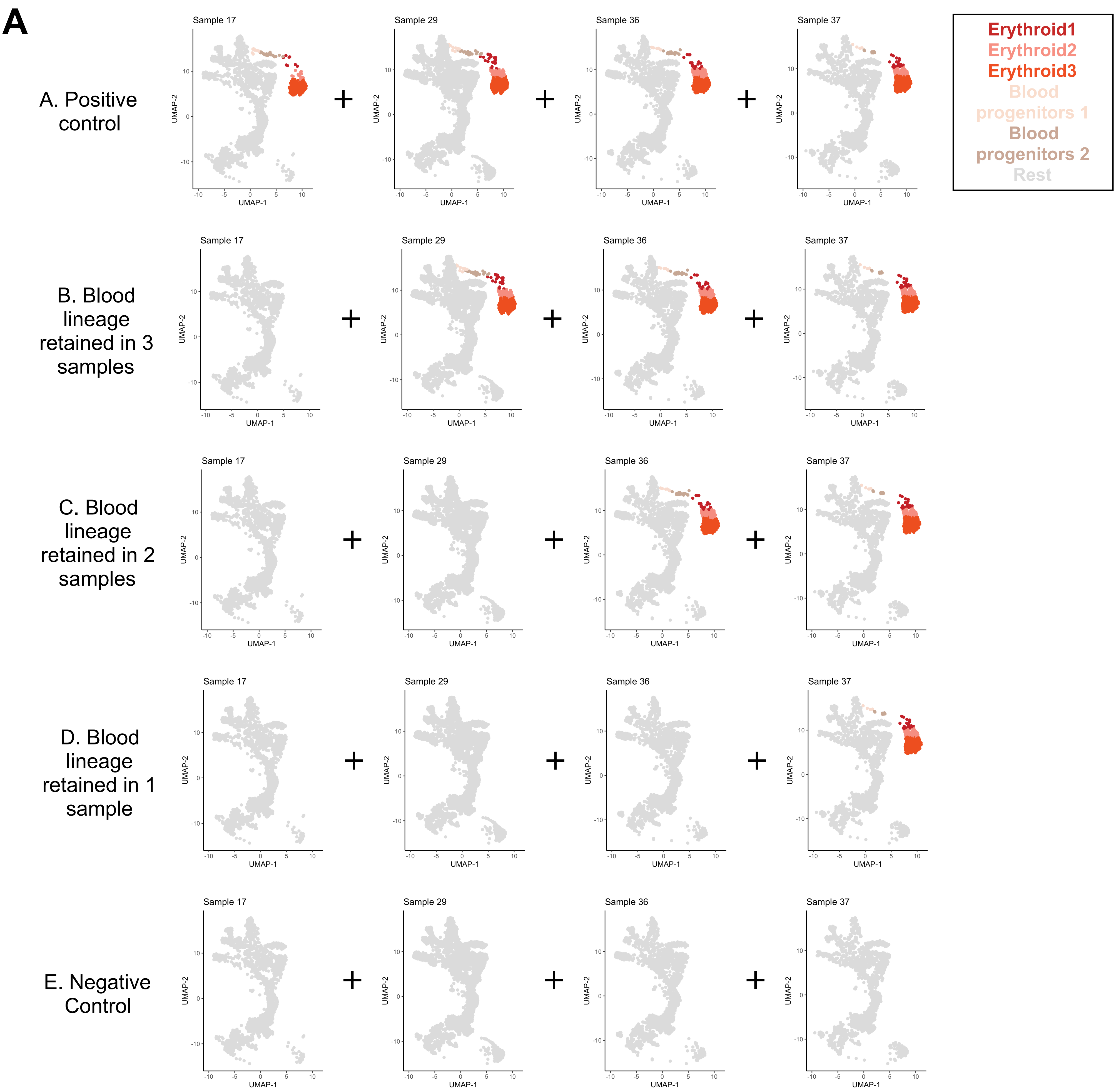


B

| | A | B | C | D | E |
|---|---|---|---|---|---|
| gene Basis | + | + | + | + | - |
| SCMER | + | + | + | - | - |

+ At least one blood marker is selected

- No blood markers are selected

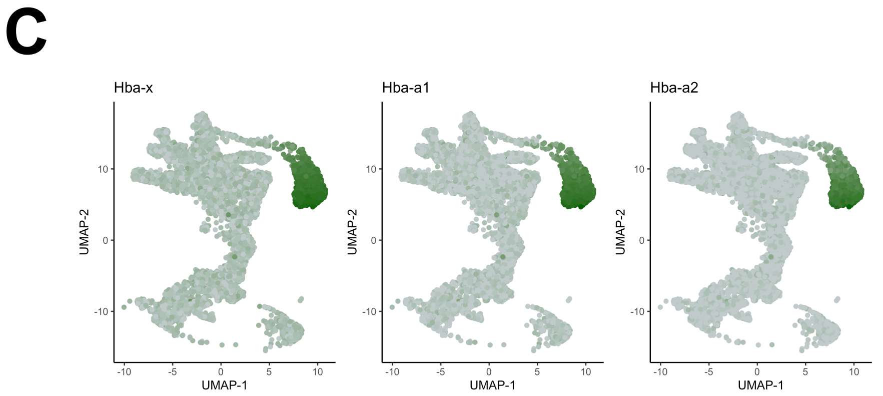

**Supplementary Figure 5. geneBasis accounts for batch effects even with highly unbalanced celltype composition.**

(A) Overview of the *in silico* experiment. Gene search was performed for 5 datasets (in rows), and inclusion of blood lineage for each sample (i.e. batch) is specified in columns.

(B) Table represents whether blood markers were selected for each dataset and method.

(C) UMAPs for mouse embryogenesis, coloured by expression of the selected blood marker genes.

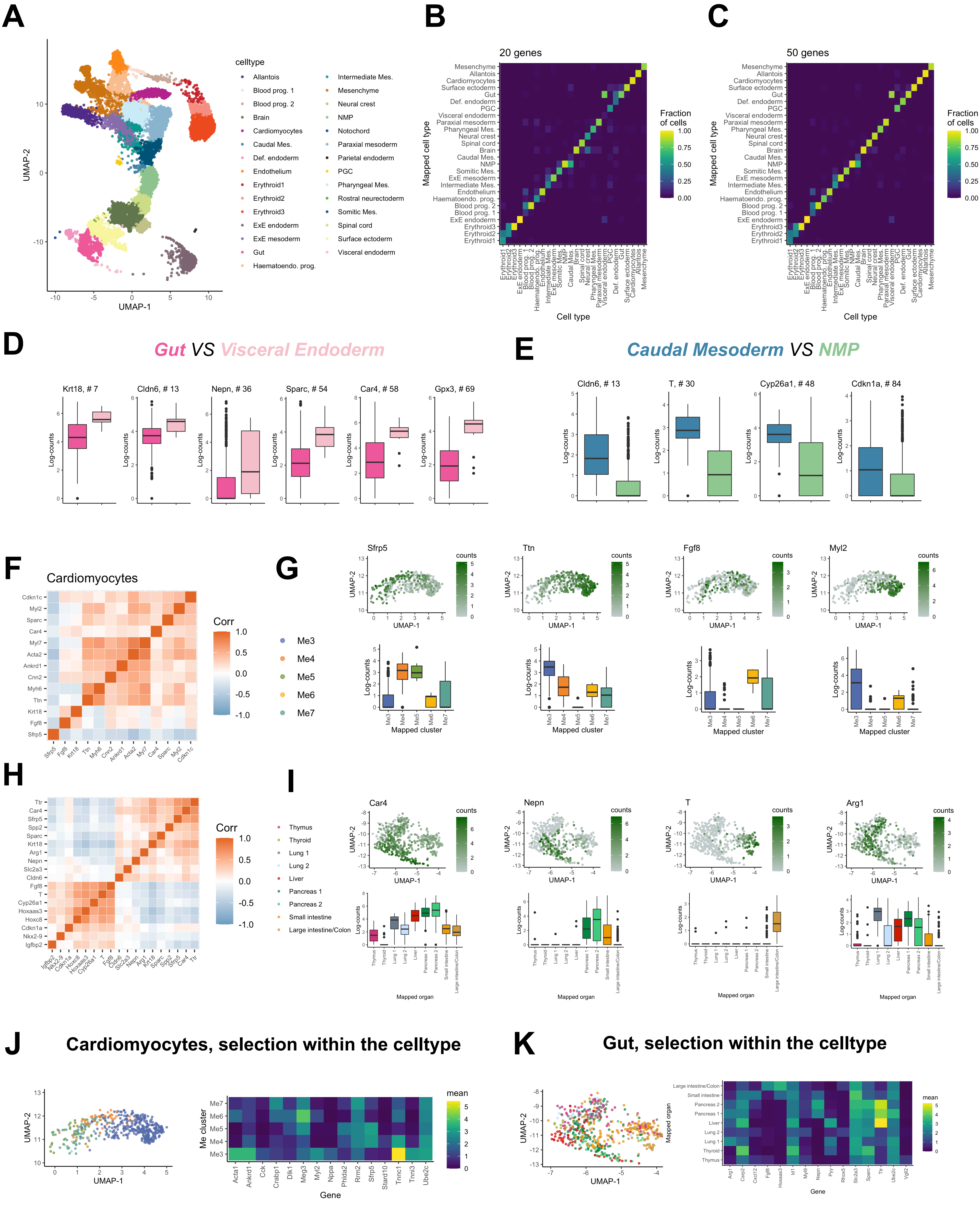
A
B
C
D
E
F
G
H
I
J
K
celltype
Allantois
Blood prog. 1
Blood prog. 2
Brain
Cardiomyocytes
Caudal Mes.
Def. endoderm
Endothelium
Erythroid1
Erythroid2
Erythroid3
ExE endoderm
ExE mesoderm
Gut
Haematoendo. prog.
Intermediate Mes.
Mesenchyme
Neural crest
NMP
Notochord
Paraxial mesoderm
Parietal endoderm
PGC
Pharyngeal Mes.
Rostral neurectoderm
Somitic Mes.
Spinal cord
Surface ectoderm
Visceral endoderm
UMAP-1
UMAP-2
20 genes
50 genes
Mapped cell type
Cell type
Fraction of cells
Gut VS Visceral Endoderm
Krt18, # 7
Cldn6, # 13
Nepn, # 36
Sparc, # 54
Car4, # 58
Gpx3, # 69
Caudal Mesoderm VS NMP
Cldn6, # 13
T, # 30
Cyp26a1, # 48
Cdkn1a, # 84
Log-counts
Cardiomyocytes
Corr
Me3
Me4
Me5
Me6
Me7
Sfrp5
Ttn
Fgf8
Myl2
counts
Mapped cluster
Thymus
Thyroid
Lung 1
Lung 2
Liver
Pancreas 1
Pancreas 2
Small intestine
Large intestine/Colon
Car4
Nepn
T
Arg1
Mapped organ
Cardiomyocytes, selection within the celltype
Gut, selection within the celltype
Me cluster
Gene
mean

**Supplementary Figure 6. Detailed analysis of the gene selection for mouse embryogenesis.**

(A) UMAP representation coloured by celltypes. Note that it duplicates part of Figure 2C and is introduced here solely to facilitate interpretation of the data.

(B) Celltype confusion matrix for the first 20 selected genes.

(C) Celltype confusion matrix for the first 50 selected genes.

(D) Box Plots representing bulk log-normalised expression across celltypes for genes that are differentially expressed between Visceral endoderm and Gut.

(E) Box Plots representing bulk log-normalised expression across celltypes for genes that are differentially expressed between Caudal mesoderm and NMP.

(F) Co-expression (within Cardiomyocytes) of genes prioritised by geneBasis and differentially expressed in Cardiomyocytes.

(G) For manually selected genes denoted as relevant to inter-cardiomyocytes heterogeneity: UMAP representation of Cardiomyocytes coloured by expression and box plots representing bulk log-normalised expression across mapped cardiac clusters.

(H) Co-expression (within Gut cells) of genes prioritised by geneBasis and differentially expressed in Gut.

(I) For manually selected genes denoted as relevant to inter-gut heterogeneity: UMAP representation of Gut cells colored by expression and box plots representing bulk log-normalised expression across mapped cardiac clusters.

(J) Left panel: UMAP representation of Cardiomyocytes, coloured by mapped cardiac clusters, when integration between datasets was performed using the first 15 genes prioritised by geneBasis if performed directly on Cardiomyocytes. Right panel: Heatmap representing average expression for assigned (using mapping with all genes) cardiac cluster (Y-axis) for first 15 genes prioritised by geneBasis if performed directly on Cardiomyocytes.

(K) Left panel: UMAP representation of Gut cells, coloured by mapped cardiac clusters, when integration between datasets was performed using first 15 genes prioritised by geneBasis if performed directly on Gut cells. Right panel: Heatmap representing average expression for assigned (using mapping with all genes) gut suborgan (Y-axis) for first 15 genes prioritised by geneBasis if performed directly on Gut cells.

#### *Spleen*

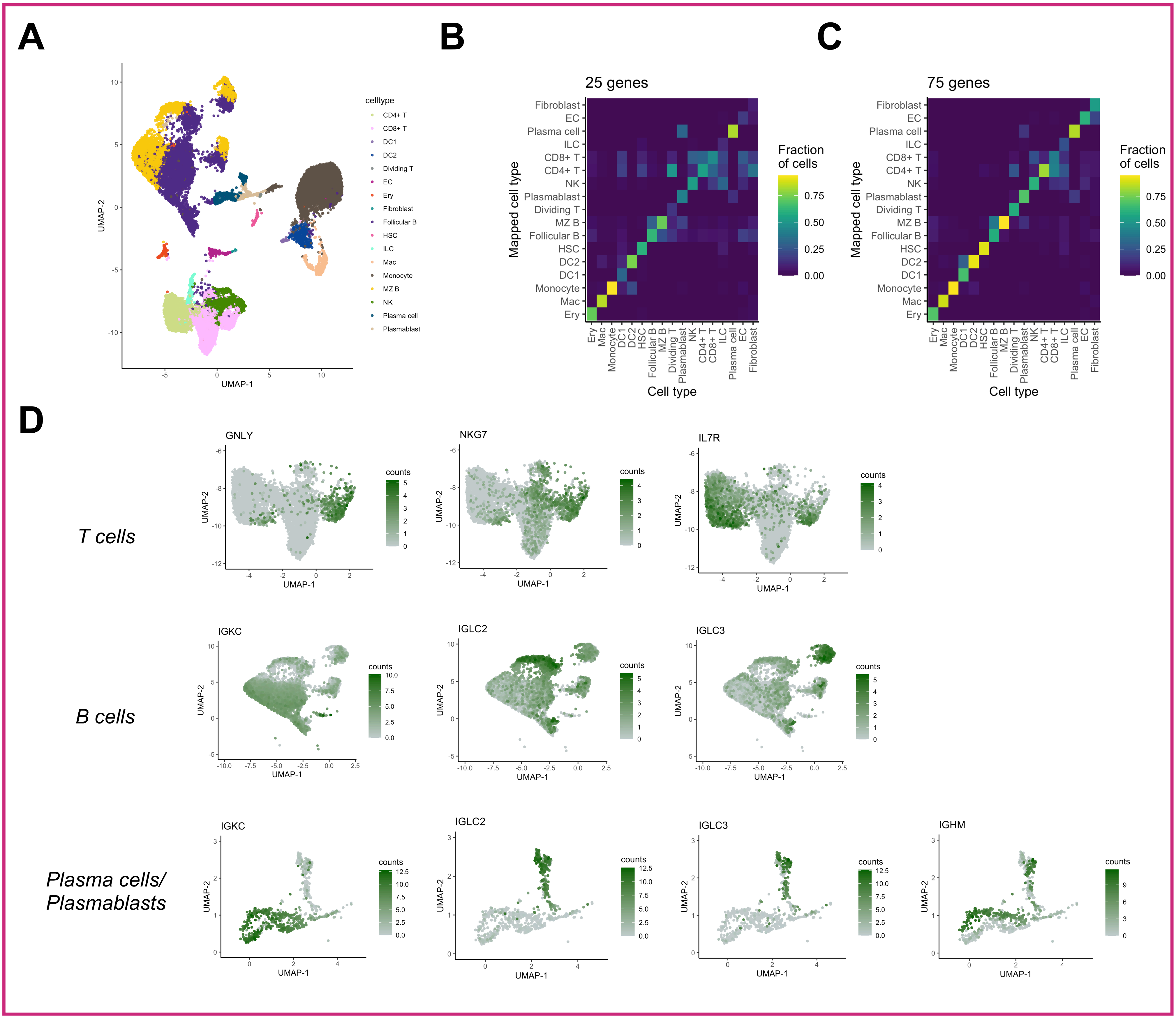


#### *Pancreas*

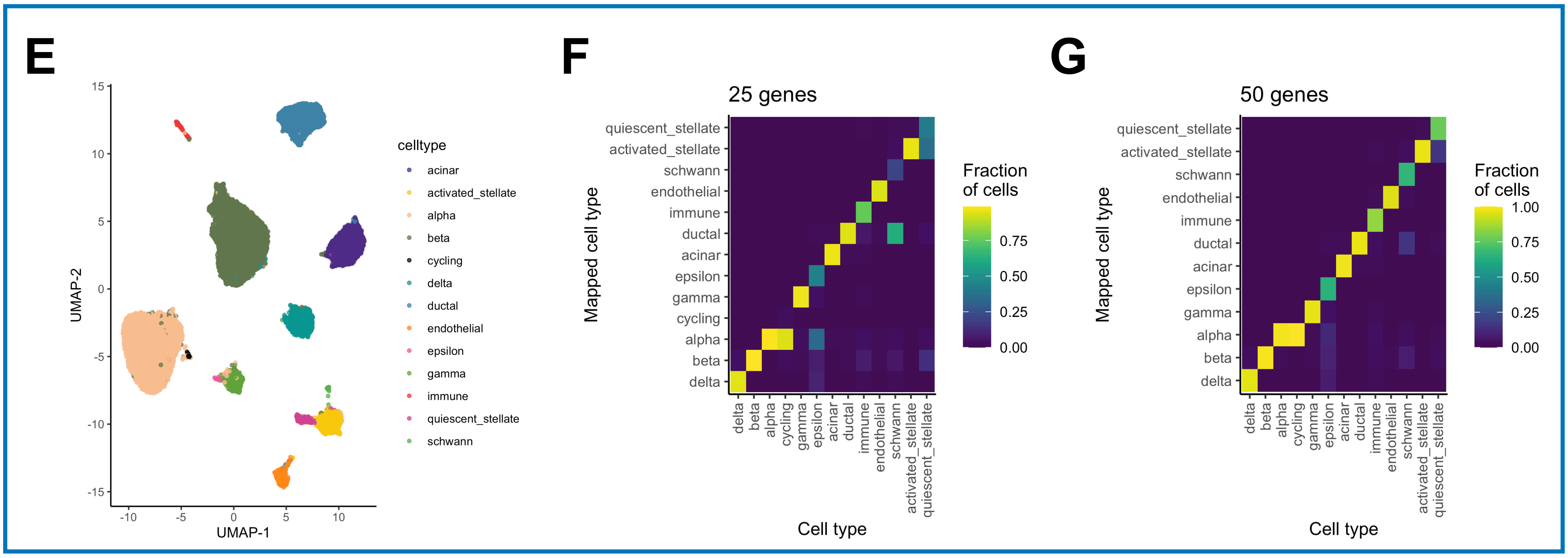

**Supplementary Figure 7. Detailed analysis of the selection for spleen and pancreas . Panels A-D correspond to spleen; panels E-G correspond to pancreas.**

(A) UMAP representation coloured by celltypes. Note that it duplicates part of Figure 2C and is introduced here solely to facilitate interpretation of the data.

(B) Celltype confusion matrix for 25 selected genes.

(C) Celltype confusion matrix for the first 75 selected genes.

(D) UMAP representation of T cells (Upper panel), B cells (Middle panel) and Plasma cells and Plamblasts (Lower panel), colored by genes relevant for inter-celltype variability within corresponding celltypes.

(E) UMAP representation colored by celltypes. Note that it duplicates part of Figure 2C and is introduced here solely to facilitate interpretation of the data.

(F) Celltype confusion matrix for 25 selected genes.

(G) Celltype confusion matrix for 50 selected genes.

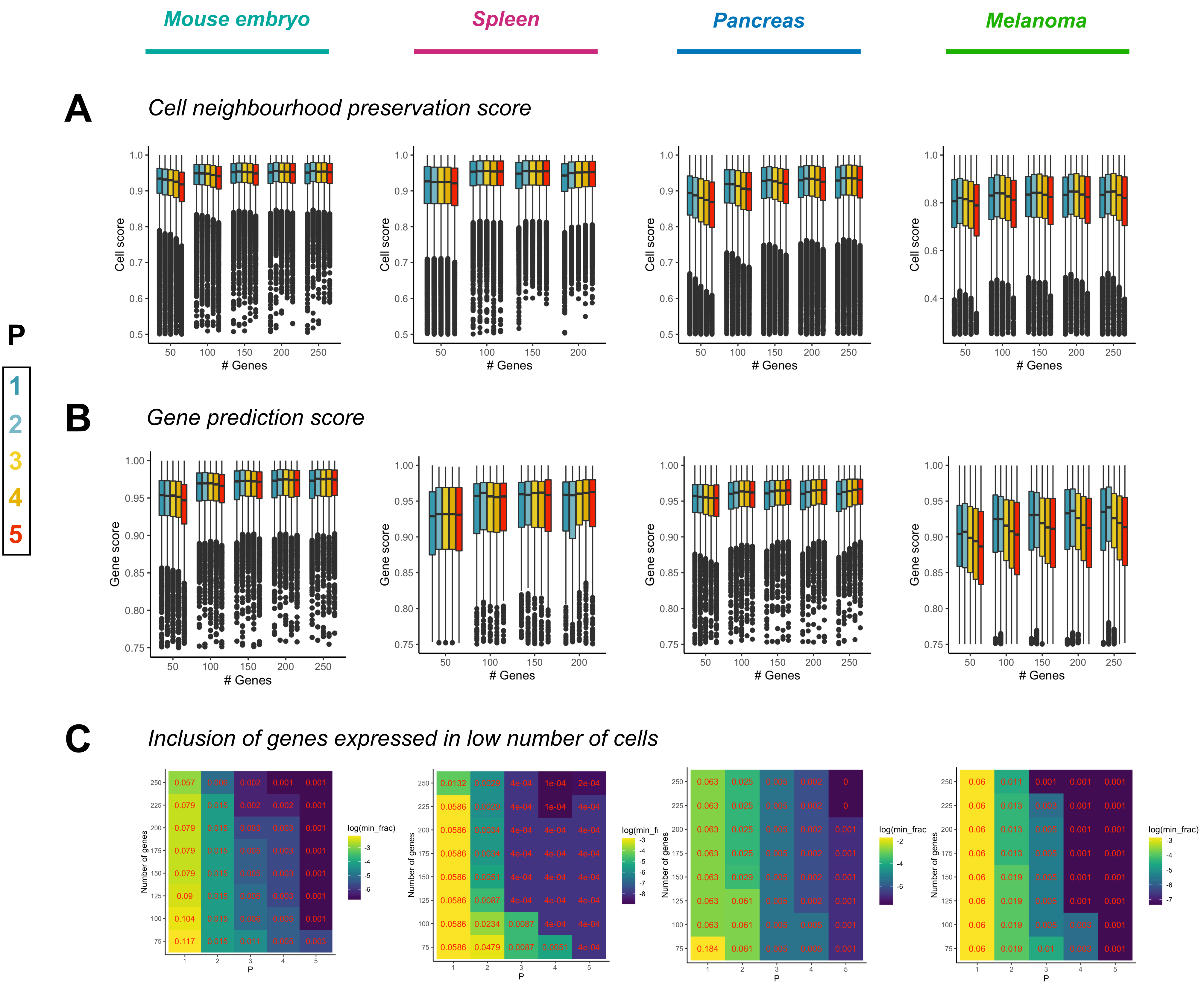

Mouse embryo
Spleen
Pancreas
Melanoma
A
Cell neighbourhood preservation score
B
Gene prediction score
C
Inclusion of genes expressed in low number of cells
P
1
2
3
4
5
Cell score
Gene score
### Genes
Number of genes
log(min_frac)

**Supplementary Figure 8. Minkowski distance of order p=3 provides the resolution to select rare markers and genes expressed in a low fraction of cells.** Each column corresponds to a single dataset.

(A) Box plots representing overall cell neighborhood preservation score distribution as a function of number of genes in the selections (X-axis) and different orders of Minkowski distance (p) used for the algorithm (in colour).

(B) Box plots representing overall gene prediction score distribution as a function of number of genes in the selections (X-axis) and p (orders of Minkowski distance) used for the algorithm (in colour).

(C) Heatmaps representing the ability of a selection with the given order of Minkowski distance to prioritise lowly expressed genes. X-axis corresponds to the choice of p, Y-axis corresponds to number of genes in the selection, and both numbers (absolute scale) and colours (log scale) correspond to the minimum of expression levels (in the context fraction of cells with non-zero counts) across selected genes.

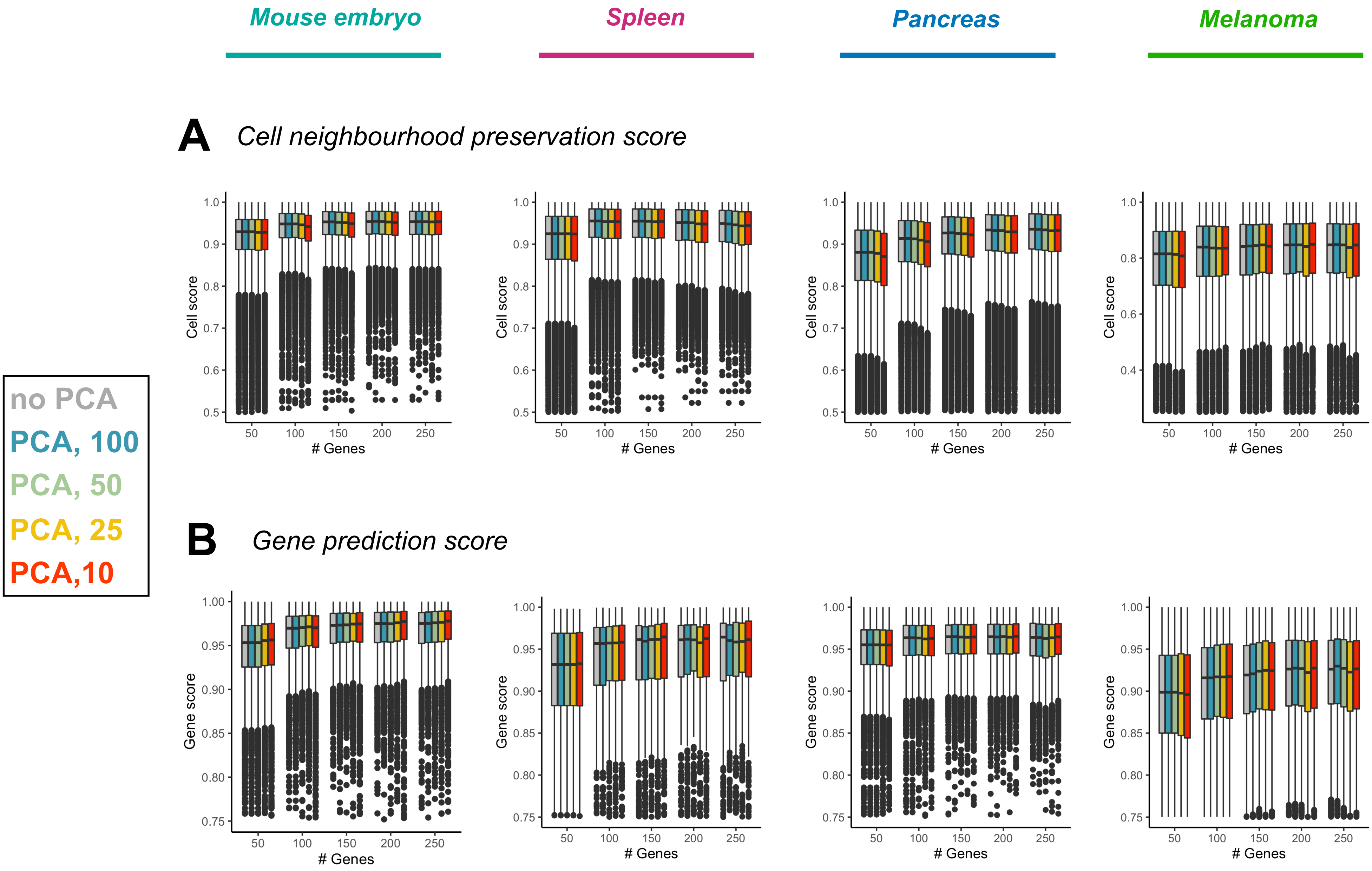

Mouse embryo
Spleen
Pancreas
Melanoma
A
Cell neighbourhood preservation score
Cell score
### Genes
50
100
150
200
250
B
Gene prediction score
Gene score
no PCA
PCA, 100
PCA, 50
PCA, 25
PCA,10

**Supplementary Figure 9.Performing PCA for mapping manifolds with selections does not change cell neighborhood preservation score or gene prediction score.**

(A) Box plots representing overall cell neighbourhood preservation score distribution as a function of the number of genes in the selections (X-axis) and optional PCA step with different number of components being selected + default no PCA (in colour).

(B) Box plots representing overall gene prediction score distribution as a function of number of genes in the selections (X-axis) and optional PCA step with different number of components being selected + default no PCA (in colour).
